# An oligomeric state-dependent switch in FICD regulates AMPylation and deAMPylation of the chaperone BiP

**DOI:** 10.1101/595835

**Authors:** Luke A. Perera, Claudia Rato, Yahui Yan, Lisa Neidhardt, Stephen H. McLaughlin, Randy J. Read, Steffen Preissler, David Ron

## Abstract

AMPylation is an inactivating modification that matches the activity of the major endoplasmic reticulum (ER) chaperone BiP to the burden of unfolded proteins. A single ER-localised Fic protein, FICD (HYPE), catalyses both AMPylation and deAMPylation of BiP. However, the basis for the switch in FICD’s activity is unknown. We report on the transition of FICD from a dimeric enzyme, that deAMPylates BiP, to a monomer with potent AMPylation activity. Mutations in the dimer interface or in residues tracing an inhibitory relay from the dimer interface to the enzyme’s active site favour BiP AMPylation in vitro and in cells. Mechanistically, monomerisation relieves a repressive effect allosterically-propagated from the dimer interface to the inhibitory Glu234, thereby permitting AMPylation-competent binding of MgATP. Whereas, a reciprocal signal propagated from the nucleotide binding site, provides a mechanism for coupling the oligomeric-state and enzymatic activity of FICD to the energy status of the ER.

**Impact Statement:** Unique amongst known chaperones, the endoplasmic reticulum (ER)-localized Hsp70, BiP, is subject to transient inactivation under conditions of low ER stress by reversible, covalent modification – AMPylation. The enzyme responsible for this modification, FICD, is in fact a bifunctional enzyme with a single active site capable of both AMPylation and deAMPylation. Here we elucidate, by biochemical, biophysical and structural means, the mechanism by which this enzyme is able to switch enzymatic modality: by regulation of its oligomeric state. The oligomeric state-dependent reciprocal regulation of FICD activity is, in turn, sensitive to the ATP/ADP ratio. This allosteric pathway potentially facilitates the sensing of unfolded protein load in the ER and permits the transduction of this signal into a post-translational buffering of ER chaperone activity.

## Introduction

In all domains of life, protein folding homeostasis is achieved by balancing the burden of unfolded proteins and the complement of chaperones. In the endoplasmic reticulum (ER) of animal cells, this match is facilitated by the unfolded protein response (UPR). In addition to well-recognized transcriptional and translational strands of the UPR (Walter & Ron, 2011), recent findings have drawn attention to the existence of rapid post-translational mechanisms that adjust the activity of the ER Hsp70 chaperone BiP. Best understood amongst these is AMPylation, the covalent addition of an AMP moiety from ATP onto a hydroxyl group-containing amino acid side chain.

AMPylation conspicuously occurs on Thr518 of BiP (Preissler *et al*, 2015b; Broncel *et al*, 2016; Casey *et al*, 2017). The resulting BiP-AMP is locked in a domain-coupled ATP-like state (Preissler *et al*, 2015b, 2017b; Wieteska *et al*, 2017). Consequently, BiP-AMP has high rates of client protein dissociation (Preissler *et al*, 2015b). Moreover, the ATPase activity of BiP-AMP is resistant to stimulation by J-protein co-factors, which greatly reduces the chaperone’s ability to form high-affinity complexes with its clients (Preissler *et al*, 2017b). AMPylation therefore serves to inactivate BiP. This modification is temporally dynamic and the levels of BiP-AMP respond to changes of the protein folding load in the ER.

Consistent with its inactivating character, BiP modification in cells is enhanced by inhibition of protein synthesis (Laitusis *et al*, 1999) or during recovery from ER stress; when BiP levels exceed the requirements of unfolded client proteins (Preissler *et al*, 2015b). Conversely, as levels of ER stress increase, modification is reversed by deAMPylation, recruiting BiP back into the chaperone cycle (Laitusis *et al*, 1999; Chambers *et al*, 2012; Preissler *et al*, 2015b). Accordingly, BiP modification creates a readily-accessible pool of latent folding capacity that buffers both ER stress (through deAMPylation) and over-chaperoning (through AMPylation). These features may contribute to the observation whereby in the *Drosophila* visual system, loss of the ability to AMPylate BiP results in light-induced blindness (Rahman *et al*, 2012; Moehlman *et al*, 2018).

AMPylation of BiP is mediated by the ER-localised enzyme FICD (filamentation induced by <u>c</u>AMP <u>d</u>omain protein, also known as HYPE) (Ham *et al*, 2014; Sanyal *et al*, 2015; Preissler *et al*, 2015b). FICD is the only known metazoan representative of a large family of bacterial Fic-domain proteins (Khater & Mohanty, 2015a). Fic proteins contain a conserved active site motif, HPFx(D/E)GN(G/K)R_1_xxR_2_, and many possess a glutamate-containing inhibitory alpha helix (α_inh_) responsible for auto-inhibition of their canonical AMPylation activity (Engel *et al*, 2012; Goepfert *et al*, 2013). FICD is a class II Fic protein (with its α_inh_ N-terminal to its Fic domain) and an ER-localised type II, single-pass transmembrane protein, with a short cytoplasmic portion and a large luminal-facing catalytic domain (Worby *et al*, 2009; Bunney *et al*, 2014).

Crystal structures of FICD and other Fic domain proteins suggest that engagement of Glu234 (of the α_inh_) with Arg374 (R_2_ of the Fic motif) prevents binding of MgATP in a conformation conducive to catalysis (Engel *et al*, 2012; Goepfert *et al*, 2013; Bunney *et al*, 2014; Truttmann *et al*, 2016). Moreover, in vitro, modification of BiP by purified FICD requires mutation of Glu234; an observation suggesting that an AMPylation repressed state is favoured by wild-type FICD. Remarkably, the Fic domain of FICD is also responsible for BiP deAMPylation; an activity that depends on Glu234 (Preissler *et al*, 2017a; Casey *et al*, 2017) and magnesium (Veyron *et al*, 2019). These findings point to deAMPylation as the default activity of the bifunctional enzyme and implicate Glu234 in a functional switch between the two antagonistic activities of the Fic active site.

The Fic domain of human FICD forms a stable back-to-back asymmetric dimer via two dimerisation surfaces (Bunney *et al*, 2014; Truttmann *et al*, 2016) and a monomerising mutation in the dimer interface of *Drosophila* FICD did not block BiP deAMPylation in vitro (Casey *et al*, 2017). Nonetheless, distantly related bacterial enzymes hint at a possible regulatory role for Fic dimerisation: a mutation in *Clostridium difficile* Fic (CdFic) dimer interface increased auto-AMPylation (Dedic *et al*, 2016) and changes in oligomeric state affected the activity of the class III Fic protein from *Neisseria meningitidis* (NmFic) (Stanger *et al*, 2016).

Here we report on the biochemical and structural basis of an oligomeric state-dependent switch in FICD’s activity, which is well suited to post-translationally regulate protein folding homeostasis in the ER.

## Results

### Disrupting the FICD dimer favours BiP AMPylation

Whilst the *FICD* gene is necessary for BiP AMPylation, over-expression of the wild-type FICD enzyme does not result in a detectable pool of BiP-AMP in cells (Preissler *et al*, 2015b). These findings were explained in terms of dominance of the deAMPylation activity of wild-type FICD, as observed in vitro (Preissler *et al*, 2017a). However, somewhere between low-level endogenous expression, which yields physiologically-regulated AMPylation, and over-expression, which precludes BiP-AMP accumulation, retrovirally-rescued *FICD*^-/-^ cells were endowed with a measure of BiP AMPylation (Figure 1A and S1A-C). This finding points to a protein-dosage effect on wild-type FICD’s activity and suggests that the enzymatic mode of (recombinant) FICD may be affected by its concentration in the ER.

**Figure 1.**
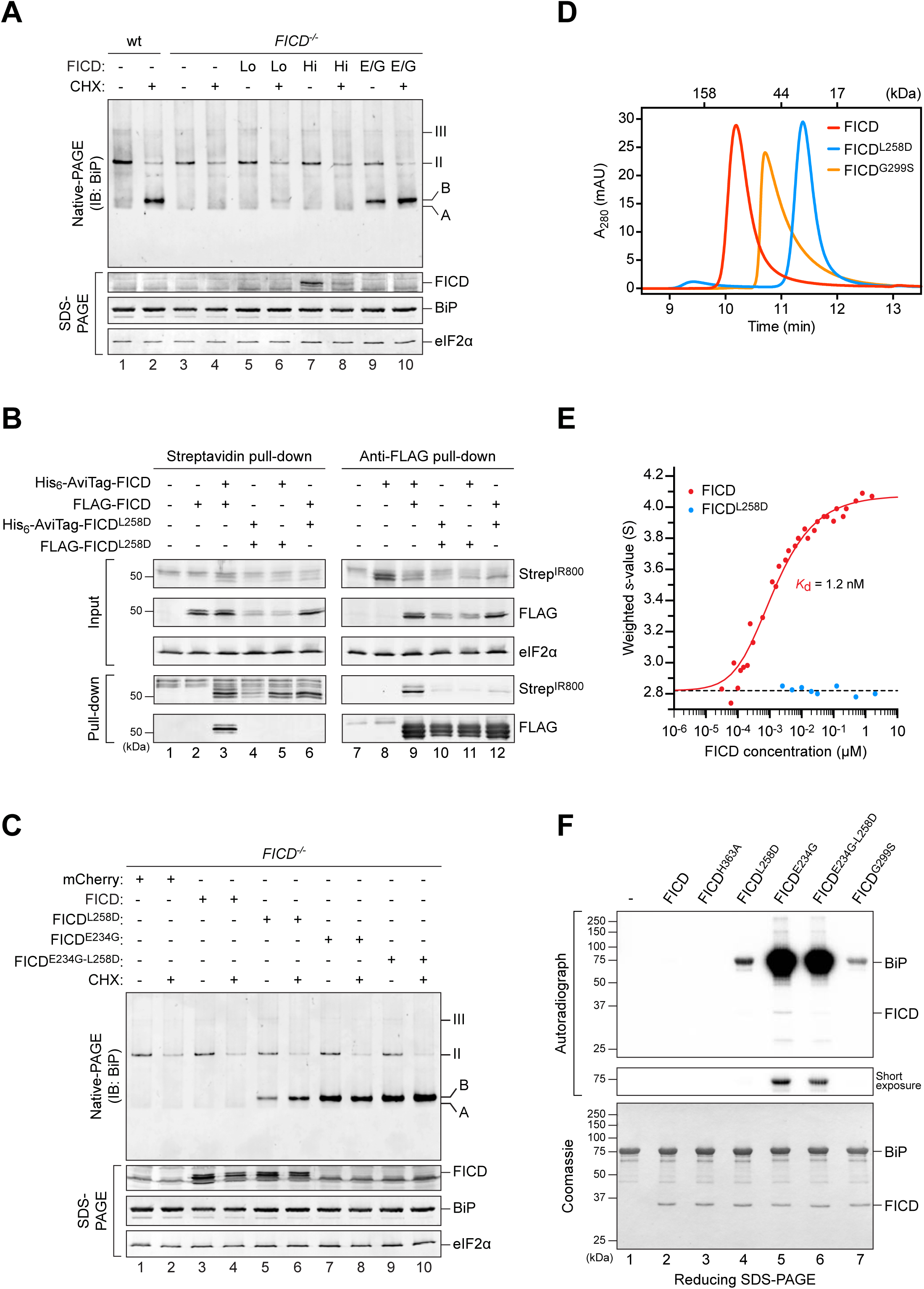
Monomeric mutant FICD promotes BiP AMPylation. A) Immunoblot of endogenous BiP resolved by native-PAGE from lysates of CHO-K1 S21 wild-type (wt) or *FICD*^-/-^ cells either transiently overexpressing wild-type FICD (high expression level; Hi) or mutant FICD^E234G^ (E/G) or stably expressing recombinant wild-type FICD (low expression level; Lo). The cells in lanes 1-4 were mock transfected. Where indicated cells were exposed to cycloheximide (CHX; 100 µg/mL) for 3 h before lysis. Unmodified (‘A’) and AMPylated (‘B’) monomeric and oligomeric (II and III) forms of BiP are indicated. Immunoblots of the same samples resolved by SDS-PAGE report on FICD, total BiP and eIF2α (loading control). Data representative of four independent experiments are shown. See Figure S1B-C.
B) Wild-type FICD forms homomeric complexes in vivo. Immunoblots of orthogonally-tagged wild-type and Leu258Asp mutant FICD in the input cell lysate and following recovery by pull-down with streptavidin (recognizing the AviTag) or anti-FLAG antibody. Proteins were detected with fluorescently-labelled streptavidin (StrepIR800) or FLAG antibody. Data representative of three independent experiments are shown.
C) Immunoblot of endogenous BiP from transfected CHO-K1 S21 *FICD*^-/-^ cells (as in *A*). Note that cells expressing monomeric FICD^L258D^ accumulate AMPylated BiP. Data representative of three independent experiments are shown.
D) Size-exclusion chromatography (SEC) analysis of wild-type and mutant FICD proteins (each at 20 µM). The elution times of protein standards are indicated as a reference. Note that the Leu258Asp mutation monomerises FICD, while Gly299Ser causes partial monomerisation. See Figure S1D-E.
E) Comparison of the signal-averaged sedimentation coefficients of wild-type (red) and monomeric mutant FICD^L258D^ (blue), as measured by analytical ultracentrifugation. A fit for monomer-dimer association (solid red line), constrained using the average value for the monomeric protein (dashed line, 2.82 S, S_w,20_ = 3.02 S), yielded a *K*_d_ of 1.2 nM with a 95% confidence interval between 1.1 to 1.4 nM and a value of 4.08 S for the dimer (S_w,20_ = 4.36 S). The fitted data points are from three independent experiments. See Figure S1F-G.
F) Autoradiograph of BiP, AMPylated in vitro by the indicated FICD derivatives, with [α-^32^P]-ATP as a substrate and resolved by SDS-PAGE. Proteins in the gel were visualized by Coomassie staining. A representative result of three independent experiments is shown.

Purified FICD forms a homodimeric complex in vitro (Bunney *et al*, 2014). Co-expression of reciprocally-tagged FICD confirmed that the wild-type protein forms homomeric complexes in cells that are disrupted by a previously characterised Leu258Asp mutation within the major dimerisation surface (Bunney *et al*, 2014) (Figure 1B). Unlike the wild-type dimerisation-competent enzyme, at a similar level of over-expression, the monomeric FICD^L258D^ yielded a clear BiP-AMP signal in *FICD*^-/-^ cells (Figure 1C). This pool was conspicuous even under basal conditions, in which wild-type cells have only a weak BiP-AMP signal, suggesting that the imposed monomeric state deregulated FICD’s activity.

Together, these observations intimate that dynamic changes in the equilibrium between the monomer and dimer may contribute to a switch between FICD’s mutually antagonistic activities – AMPylation and deAMPylation of BiP. Increasing its concentration by over-expression favours FICD dimerisation and thus perturbs such regulatory transitions. This could account for the observation that FICD overexpression, in unstressed wild-type cells, abolishes the small pool of BiP-AMP normally observed under basal conditions (Preissler *et al*, 2017b).

Size-exclusion chromatography (SEC) and analytical ultracentrifugation (AUC), with purified proteins, confirmed the stability of the FICD dimer (Figure 1D-E and S1D-G). These techniques also confirmed the strong disrupting effect of the Leu258Asp mutation (in the principal dimer surface) and revealed a weaker disrupting effect of a Gly299Ser mutation (in the secondary dimer surface) (Figure S1D-G). AUC yielded a 1.2 nM dimer dissociation constant (*K*_d_) of wild-type FICD and SEC indicated a *K*_d_ in the millimolar range for FICD^L258D^ and a *K*_d_ of 9.5 µM for FICD^G299S^. We therefore conclude that between 0.2 µM and 5 µM (concentrations at which the experiments that follow were performed) the wild-type protein is dimeric, FICD^L258D^ is monomeric, and FICD^G299S^ is partially monomeric.

In the presence of [α-^32^P]-ATP both FICD^L258D^ and FICD^G299S^ established a pool of AMPylated, radioactive BiP in vitro [Figure 1F; also observed in the *Drosophila* counterpart of FICD^L258D^ (Casey *et al*, 2017)], whereas the wild-type enzyme did not, as previously observed (Preissler *et al*, 2015b, 2017a). BiP is a substrate for AMPylation in its monomeric, ATP-bound, domain-docked conformation (Preissler *et al*, 2015b, 2017b). These experiments were therefore performed with an ATPase-deficient, oligomerisation-defective, ATP-bound BiP mutant, BiP^T229A-V461F^. Thus, the BiP-AMP signal is a result of the concentration of substrate (unmodified and modified BiP) and the relative AMPylation and deAMPylation activities of the FICD enzyme. As expected, a strong BiP-AMP signal was elicited by the unrestrained AMPylation-active FICD^E234G^ (which cannot deAMPylate BiP). FICD^E234G-L258D^ gave rise to a similar, but reproducibly slightly weaker, BiP-AMP signal relative to FICD^E234G^.

### Monomerisation switches FICD’s enzymatic activities

The ability of the dimer interface FICD mutants to yield a detectable BiP-AMP signal in vitro agreed with the in vivo data and suggested a substantial change in the regulation of the enzyme’s antagonistic activities – either inhibition of deAMPylation, de-repression of AMPylation, or a combination of both. To distinguish between these possibilities, we analysed the deAMPylation activities of the FICD mutants in an assay that uncouples deAMPylation from AMPylation. As previously observed, wild-type FICD caused the release of fluorescently labelled AMP from in vitro AMPylated BiP, whereas FICD^E234G^ did not (Preissler *et al*, 2017a) (Figure 2A). FICD^L258D^ and FICD^G299S^ consistently deAMPylated BiP 2-fold slower than the wild-type (Figure 2A and S2A). The residual in vitro deAMPylation activity of FICD^L258D^ and the absence of such activity in FICD^E234G^ is consistent with the divergent effect of expressing these deregulated mutants on a UPR reporter in cells (Figure S2B-C).

**Figure 2.**
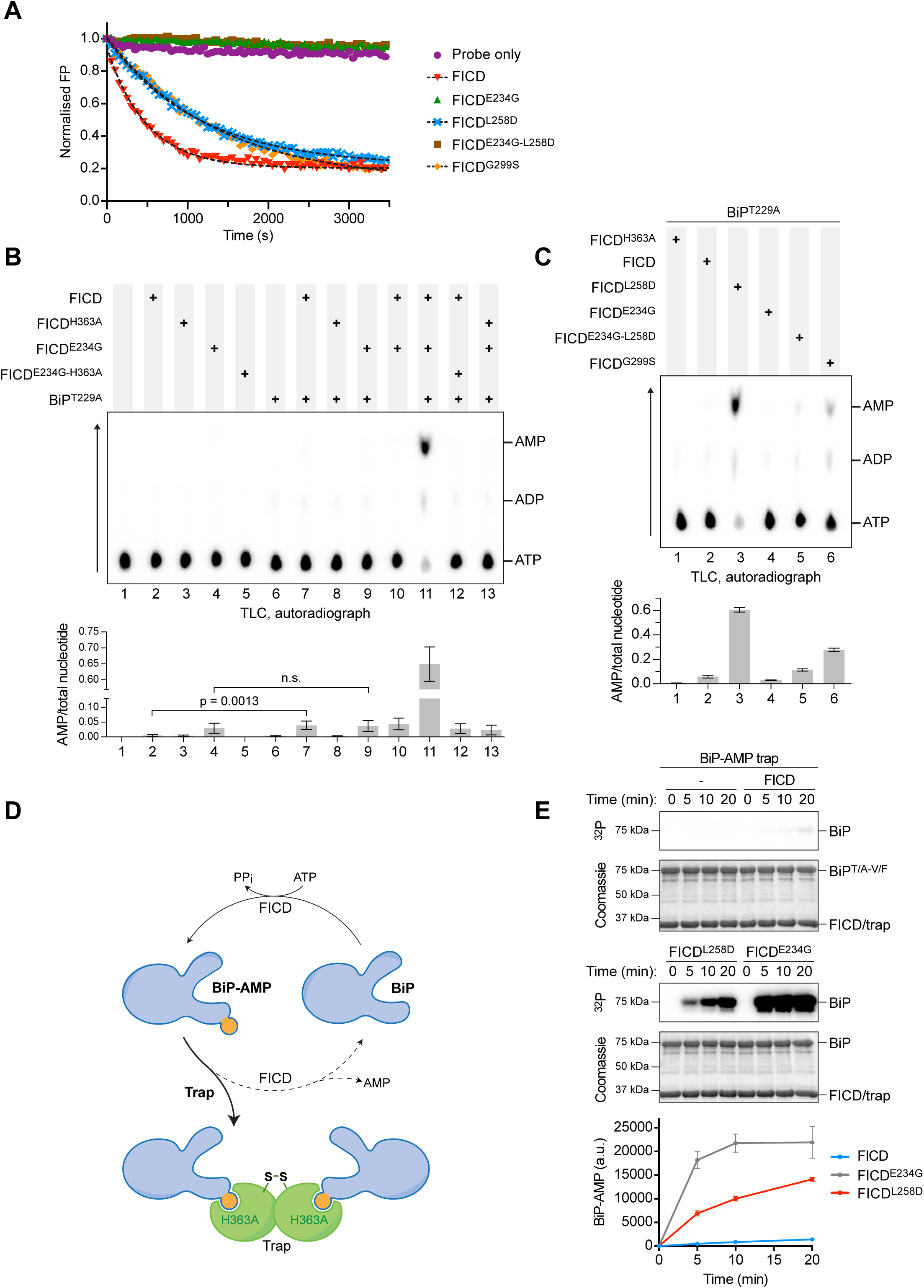
Monomerising mutations de-repress FICD’s AMPylation activity. A) Monomerising FICD mutations inhibit deAMPylation. Shown is a representative plot of data points and fit curves of the time-dependent deAMPylation of a fluorescent BiP^V461F^-AMP^FAM^ by the indicated FICD proteins (at 7.5 µM) as detected by a change in fluorescence polarisation (FP). DeAMPylation rates calculated from independent experiments are given in Figure S2A.
B-C) Dimer interface mutants both AMPylate and deAMPylate BiP. Shown are representative autoradiographs of thin layer chromatography (TLC) plates revealing AMP produced from reactions containing [α-^32^P]-ATP and the indicated FICD enzymes in the presence or absence of the co-substrate BiP (arrow indicates direction of nucleotide migration). The radioactive signals were quantified and the AMP signals were normalised to the total nucleotide signal in each sample. Plotted below are mean values ± SD from at least three independent experiments. Unpaired t-tests were performed. See Figure S2D.
D) Cartoon depicting sequestration of AMPylated BiP by a covalently linked, disulphide-stapled, _S-S_FICD^A252C-H363A-C421S^ dimer (trap). See Figure S2E-H.
E) Detection of the time-dependent accumulation of AMPylated BiP^T229A-V461F^ in radioactive reactions, containing [α-^32^P]-ATP and the indicated FICD proteins, in the presence of excess trap. At the specified time-points samples were taken and analysed by SDS-PAGE. The autoradiograph (^32^P) illustrates the radioactive signals, which represent AMPylated BiP; proteins were visualized by Coomassie staining. The radioactive signals were quantified and presented in the graph below. Mean values ± SD of three independent experiments are shown.

The FICD-mediated BiP AMPylation/deAMPylation cycle converts the co-substrate ATP to the end products AMP and pyrophosphate (Preissler *et al*, 2017a). We exploited this feature to quantify enzymatic activity. FICD was incubated with [α-^32^P]-ATP, either in the presence or absence of ATPase-deficient BiP^T229A^ and accumulation of radioactive AMP was measured by thin layer chromatography. Only background levels of AMP were generated by catalytically inactive FICD^H363A^ or FICD^E234G-H363A^ (Figure 2B). The deregulated, deAMPylation-defective FICD^E234G^ yielded a weak AMP signal that was not increased further by the presence of BiP, suggesting that the Glu234Gly mutation enables some BiP-independent ATP hydrolysis to AMP. Conversely, small but significant amounts of AMP were produced by wild-type FICD but in a strictly BiP-dependent fashion (Figure 2B-C and Figure S2D). These observations are consistent with a slow, FICD-driven progression through the BiP AMPylation/deAMPylation cycle indicating incomplete repression of wild-type FICD’s AMPylation activity under these conditions. As expected, abundant BiP-dependent AMP production was observed in reactions containing AMPylation-active FICD^E234G^ alongside deAMPylation-active wild-type FICD (Figure 2B, lane 11). Importantly, large amounts of AMP were also generated when BiP was exposed to FICD^L258D^ and, to lesser extent, FICD^G299S^ (Figure 2C and S2D). Together, these observations suggest that the AMPylation activities of the monomeric FICD mutants are significantly enhanced relative to the wild-type, whilst their deAMPylation activities are more modestly impaired.

To directly assess the AMPylation activities of bifunctional FICDs we exploited the high affinity of the catalytically inactive FICD^H363A^ for BiP-AMP, as a “trap” that protects BiP-AMP from deAMPylation (Figure 2D). To disfavour interference with the FICD enzyme being assayed we engineered the trap as a covalent disulfide linked dimer incapable of exchanging subunits with the active FICD being assayed. A cysteine (Ala252Cys) was introduced into the major dimerisation surface of the trap. To preclude aberrant disulphide bond formation, the single endogenous cysteine of FICD was also replaced (Cys421Ser). After purification and oxidation, this protein (_S-S_FICD^A252C-H363A-C421S^; the trap) formed a stable disulphide-bonded dimer (Figure S2E-F) that tightly bound BiP-AMP with fast association and slow dissociation kinetics (Figure S2G-H). Moreover, the binding of the trap to unmodified BiP was, in comparison, negligible (Figure S2G). We reasoned that adding the trap in excess to reactions assembled with BiP, ATP and FICD would sequester the BiP-AMP product and prevent its deAMPylation, enabling the comparison of AMPylation rates in isolation from the deAMPylating activity.

In presence of the trap, wild-type FICD produced a detectable BiP-AMP signal; but not in the absence of the trap (compare Figures 1F and 2E). Importantly, presence of the trap revealed that AMPylation of BiP was greatly accelerated by FICD monomerisation (> 19-fold compared to the wild-type) (Figure 2E). As expected, BiP AMPylation by FICD^E234G^ was even faster.

If the enhanced AMPylation activity of the dimerisation-defective mutants, observed above, truly represents divergent enzymatic activities of different FICD oligomeric states, it should be possible to reveal this feature by diluting the wild-type enzyme to concentrations at which an appreciable pool of monomer emerges. In AMPylation reactions set up with [α-^32^P]-ATP a detectable signal from radiolabelled BiP-AMP was noted at enzyme concentrations near the *K*_d_ of dimerisation (between 10 and 2.5 nM; Figure 3A, left). The inverse relationship of enzyme concentration to the BiP-AMP signal likely reflects the opposing activities and relative populations of AMPylation-biased FICD monomers and the deAMPylation-biased FICD dimers in each reaction. This counter-intuitive relationship of enzyme to product is resolved in the presence of the AMPylation trap; the BiP-AMP signal increased in a time-and enzyme concentration-dependent manner, as expected from a reaction which is proportional to the absolute concentration of monomeric enzyme (Figure 3A, right). In the presence of the trap the shift in the peak of the BiP-AMP signal, after 16 hours, towards lower concentrations of FICD, likely reflects incomplete protection of AMPylated BiP by the trap and its enhanced susceptibility to deAMPylation at higher concentrations of (dimeric) FICD.

**Figure 3.**
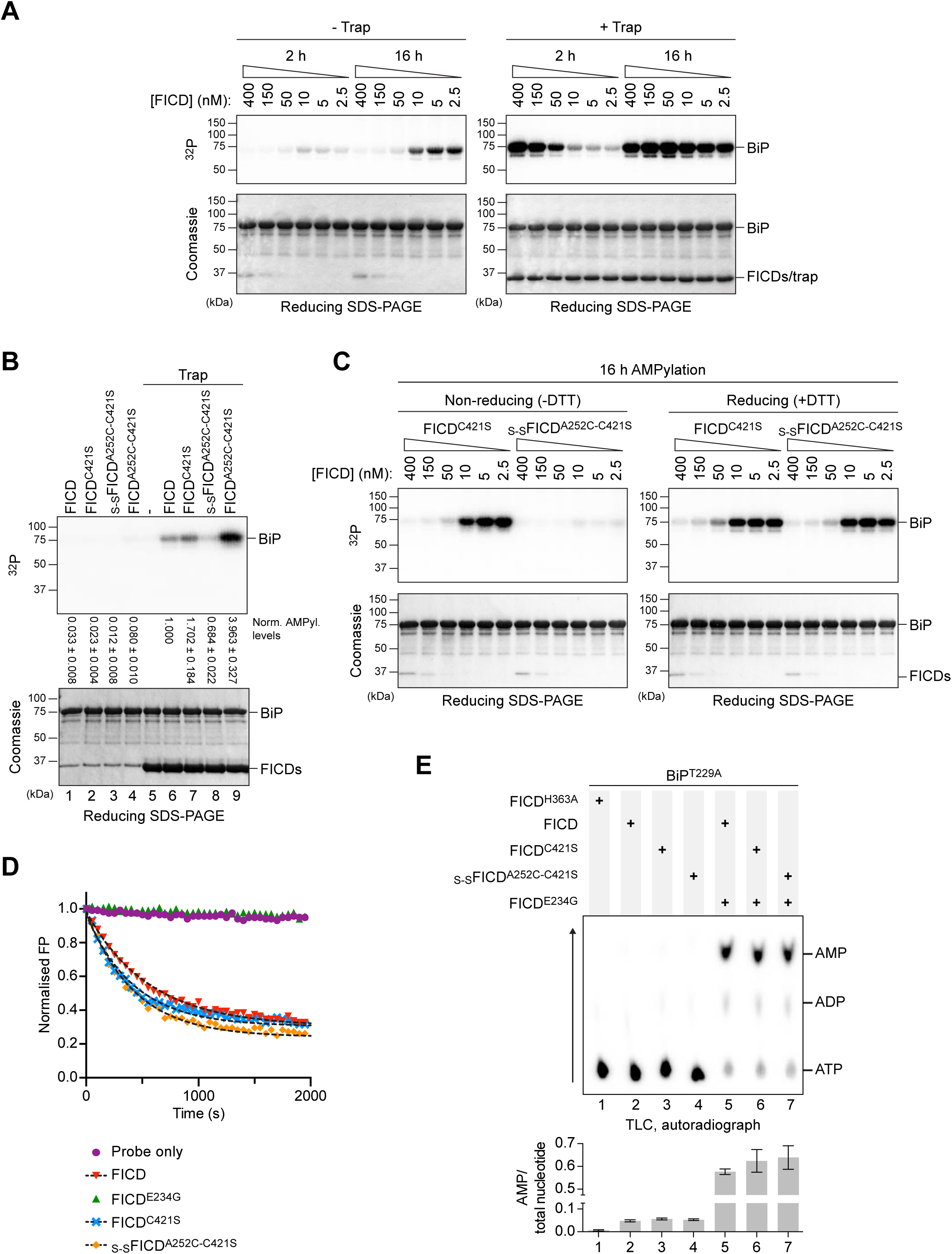
Monomerisation by dilution enhances the AMPylation activity of wild-type FICD. A) Autoradiographs of in vitro reactions containing varying concentration of wild-type FICD protein and fixed concentrations of BiP^T229A-V461F^ and [α-^32^P]-ATP as co-substrates, resolved by SDS-PAGE after the indicated incubation times. The proteins were visualized by Coomassie staining of the gel (bottom). The reactions shown on the right were performed in the presence of an excess of _S-S_FICD^A252C-H363A-C421S^ (trap) to delay de-modification of BiP. Representative gels are shown, and similar results were observed in three independent experiments.
B) As in *(A)* but with 0.2 µM of the indicated FICD variant. The radioactive signals were detected by autoradiography, quantified, and normalised to the signal in lane 6. The mean radioactive signals ± SD from three independent experiments are given. The proteins were visualized by staining with Coomassie. See Figure S3A-B.
C) As in *(A)* but with dilutions of FICD^C421S^ or covalently linked _S-S_FICD^A252C-C421S^. Reactions were preceded by a 16 h incubation of FICD in presence or absence of the reducing agent (DTT). Representative gels are shown of three independent experiments. See Figure S3C.
D) Forced dimerisation does not significantly alter deAMPylation rates. Time-dependent deAMPylation of fluorescent BiP^V461F^-AMP^FAM^ by the indicated FICD proteins (at 7.5 µM) assayed by fluorescence polarisation (as in Figure 2A). A representative experiment (data points and fit curves) is shown and rates are given in Figure S2A. See Figure S3D.
E) Representative autoradiograph of thin layer chromatography (TLC) plates revealing AMP produced from reactions containing [α-^32^P]-ATP and the indicated FICD enzymes in the presence of the co-substrate BiP. AMP signals were normalised to the total nucleotide signal in each sample and the graph below plots mean values ± SD from at least three independent experiments.

If monomerisation significantly enhances AMPylation activity, constitutive FICD dimers that are unable to dissociate should have low AMPylation activity and fail to produce modified BiP even under dilute conditions. To test this prediction, we created a disulphide-linked wild-type FICD (_S-S_FICD^A252C-C421S^), which, after purification and oxidation, formed a covalent dimer (Figure S3A). Moreover, its SEC profile was indistinguishable from wild-type FICD or the cysteine-free counterpart, FICD^C421S^ (Figure S3B). In the presence of the BiP-AMP trap, oxidised _S-S_FICD^A252C-C421S^ produced significantly less AMPylated BiP than either wild-type or FICD^C421S^ at similar concentrations (Figure 3B, lane 8 and S3C).

Repression of AMPylation was imposed specifically by the covalent dimer, as non-oxidised FICD^A252C-C421S^ elicited a conspicuous pool of BiP-AMP - more than the wild-type enzyme (Figure 3B, lane 9 and S3C) - an observation explained by the weakening of the FICD dimer imposed by the Ala252Cys mutation (Figure S1D-E). Similarly, in absence of the trap, the ability of pre-oxidised _S-S_FICD^A252C-C421S^ to establish a pool of AMPylated BiP was greatly enhanced by diluting the enzyme into a buffer containing DTT. FICD^C421S^, by contrast, produced similar amounts of modified BiP under both non-reducing and reducing conditions (Figure 3C).

DeAMPylation activities of oxidised and non-oxidised FICD^A252C-C421S^ were comparable and similar to wild-type FICD (Figure 3D-E, S2A and S3D), pointing to the integrity of these mutant enzymes. Together, these observations argue that covalent _S-S_FICD^A252C-C421S^ dimers selectively report on the enzymatic characteristics of wild-type FICD in its dimeric state. This protein therefore serves to help validate the conclusion that a low concentration of wild-type FICD favours the formation of monomers, whose AMPylation activity is de-repressed, to promote BiP modification.

### An AMPylation-repressive signal is transmitted from the dimer interface to the active site

The crystal structure of dimeric FICD suggests the existence of a hydrogen-bond network, involving the side-chains of Lys256 and Glu242, linking the dimer interface with the enzyme’s active site, impinging on the AMPylation-inhibiting Glu234 (Figure 4A). To test this notion, we mutated both putative dimer relay residues. FICD^K256S^ and FICD^E242A^ formed stable dimers, as assayed by SEC, with dimer *K*_d_ values under 400 nM (Figure 4B and S1D-E). In vitro both mutants established a pool of modified BiP (Figure 4C and S4A). This remained the case even at FICD concentrations in which negligible amounts of monomer are predicted (2 and 10 µM; Figure S4A). De-repression of AMPylation by these dimer relay mutations was also evidenced by the enhanced BiP-dependent AMP production, relative to wild-type FICD (Figure 4D), whilst deAMPylation activities were similar (Figure S2A and S4B). Combining the Lys256Ser and the monomerising Leu258Asp mutations (FICD^K256S-L258D^) further enhanced the BiP-AMP pool produced in vitro (Figure S4A), an observation only partially attributable to the concomitant decrease in the deAMPylation rate (Figure S2A and S4C). These observations suggest that residues connecting the dimer interface and the active site contribute to repression of AMPylation and that mutating these residues uncouples a gain-of-AMPylation activity from the oligomeric state of FICD.

**Figure 4.**
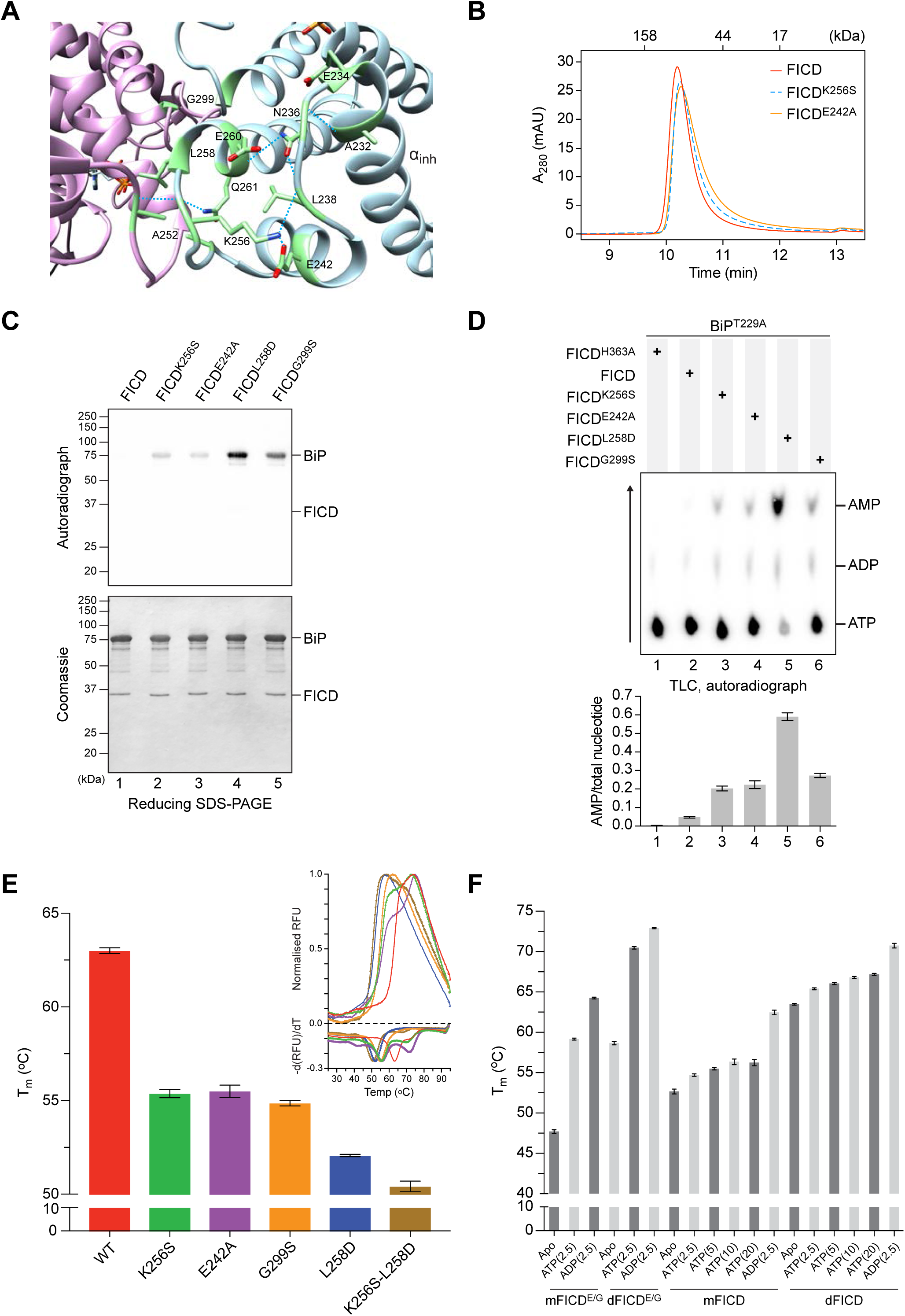
Residues connecting the FICD dimer interface with the inhibitory ɑ-helix stabilise FICD and repress AMPylation. A) Ribbon diagram of the FICD dimer interface with monomers in purple and blue ribbons (PDB:6i7g). Residues involved in a H-bond network linking the dimer interface to the α_inh_ (as well as Gly299 and Glu234) are shown as green sticks. Sub-3.50 Å hydrogen bonds made by Asn236, Leu238 and Lys256 are depicted as dotted cyan lines.
B) Size-exclusion chromatography elution profile of wild-type and mutant FICD proteins (each at 20 µM). Protein absorbance at 280 nm is plotted against elution time. The elution times of protein standards are indicated as a reference.
C) Radioactive in vitro AMPylation reactions containing the indicated FICD proteins, [α-^32^P]-ATP, and BiP^T229A-V461F^ were analysed by SDS-PAGE. The radioactive BiP-AMP signals were detected by autoradiography and proteins were visualized by Coomassie staining of the gel. See Figure S4A.
D) Representative autoradiograph of thin layer chromatography (TLC) plates revealing AMP produced from reactions containing [α-^32^P]-ATP and the indicated FICD enzymes in the presence of the co-substrate BiP. The radioactive signals were quantified and the AMP signals were normalised to the total nucleotide signal in each sample. The graph shows mean AMP values ± SD from three independent experiments.
E) Melting temperatures (T_m_) of the indicated FICD mutants (at 2 µM) were measured by differential scanning fluorimetry (DSF). Shown is the mean T_m_ ± SD of three independent experiments. The inset shows melt curves with their negative first derivatives from a representative experiment. See Figure S4D.
F) A plot of the melting temperature of the indicated FICD proteins in absence (Apo) or presence of nucleotides. Shown are the mean T_m_ values ± SD of three independent DSF experiments. Monomeric FICD^L258D^ (mFICD) and FICD^L258D-E234G^ (mFICD^E/G^) as well as dimeric wild-type FICD (dFICD) and FICD^E234G^ (dFICD^E/G^) were tested. ADP and ATP concentrations in mM are given in parentheses. See Figure S4E for *K*_½_ quantification.

Transmission of a repressive signal via a network of intramolecular interactions is also supported by the correlation between de-repression of BiP AMPylation and the negative effect of various mutants on the global stability of FICD. Differential scanning fluorimetry (DSF) revealed an inverse relationship between the AMPylation activity and the melting temperature (T_m_) of FICD mutants (Figure 4E and S4D). These differences in flexibility were observed despite the fact that the DSF assays were conducted at relatively high protein concentrations (2 µM) that would favour dimerisation of all but the most dimerisation-defective mutants.

Nucleotide binding stabilises all FICD variants (Figure S4D), a feature that is conspicuous in case of the AMPylation de-repressed FICD^E234G^ (Bunney *et al*, 2014). However, monomerisation imposed by the Leu258Asp mutation, did not significantly increase ATP-induced stabilisation of FICD (ΔT_m_) (Figure 4F and S4E). Interestingly, although AMPylation activity correlated with increased FICD flexibility this was not reflected in an appreciably altered propensity to bind ATP. This suggested that the variation in enzyme activity of different FICD mutants may arise not from variation in their affinity for nucleotide but from their particular mode of ATP binding. To explore this possibility, we set out to co-crystallise FICD variants with MgATP.

### Monomerisation favours AMPylation-competent binding of MgATP

High-resolution X-ray crystal structures of monomeric and dimeric FICD were obtained in various nucleotide bound states (Table 1). The tertiary structure of the Fic domain of both the monomeric FICD^L258D^ and the dimeric relay mutant FICD^K256S^ deviated little from that of the nucleotide-free wild-type dimer structure (FICD:Apo; PDB: 4u04) (Figure 5A and S5A). Moreover, co-crystallisation of FICD^L258D^, FICD^K256A^ or the wild-type dimer with ATP or an ATP analogue (AMPPNP) also resulted in no significant Fic domain conformational change from FICD:Apo (Figure 5A and S5A). Accordingly, the greatest root-mean squared deviation (RMSD) between the Fic domain of the FICD:ATP structure and any other monomeric or dimer relay FICD structure is 0.53 Å (observed between FICD:ATP and FICD^L258D^:Apo; residues 213-407). The only conspicuous change in global tertiary structure occurred in the TPR domain of FICD^L258D^ co-crystallised with ATP or AMPPNP, in which the TPR domain is flipped almost 180° from its position in other FICD structures (Figure 5A). Notably, in all FICD structures the α_inh_ remains firmly juxtaposed to the core Fic domain.

**Figure 5.**
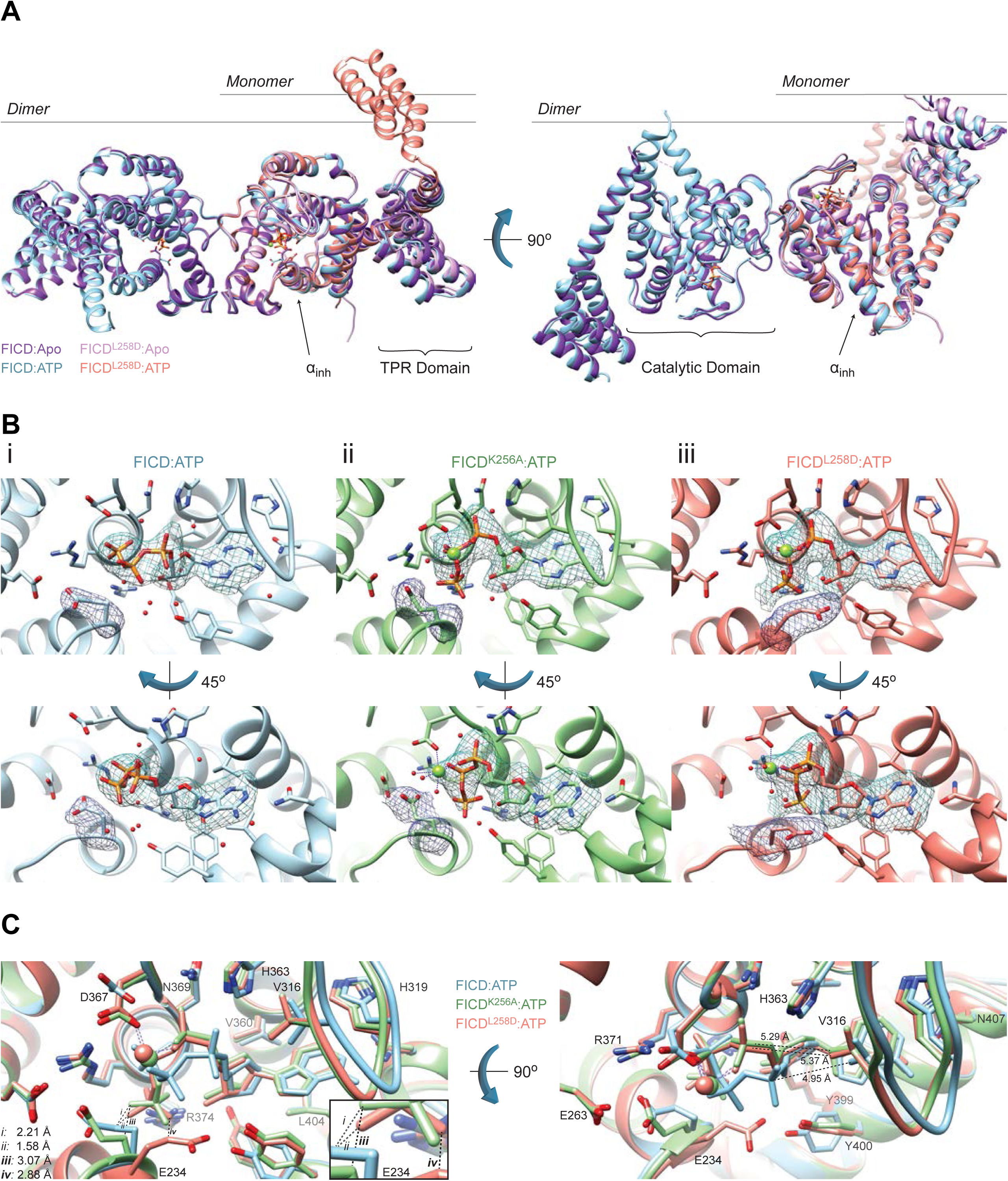
Monomeric FICD binds ATP in an AMPylation-competent conformation. A) Monomerisation does not result in large conformational changes in FICD. Shown is the alignment, from residues 213-407, of FICD molecules in the asymmetric unit. Monomeric FICD^L258D^ and dimeric wild-type FICD ± ATP, are coloured as indicated. Glu234, ATP (and Mg, where applicable), are shown as sticks (or green spheres). The inhibitory alpha helix (α_inh_) and gross domain architecture is annotated. Note the only significant deviation in tertiary structure is the flipping of the TPR domain in the FICD^L258D^:ATP structure. The FICD:Apo structure is from PDB: 4U0U. See Figure S5A.
B) Cocrystallisation of FICD variants with MgATP results in electron densities for nucleotide and the inhibitory Glu234. Unbiased polder (OMIT) maps for ATP (± Mg) and Glu234 are shown as blue and purple meshes, respectively. ***(i)*** The wild-type dimer FICD structure displays a lack of density corresponding to a Mg^2+^ ion. The ATP density is contoured at 3.5 σ and the Glu234 at 5.0 σ. ***(ii)*** The dimeric dimer relay mutant FICD^K256A^ displays a clear MgATP density up to and including the γ-phosphate phosphorous atom. The ATP density and Glu234 densities are both contoured at 3.0 σ. ***(iii)*** Monomeric FICD^L258D^ shows a clear MgATP density. The ATP density is contoured at 3.0 σ and the Glu234 at 5.0 σ. All residues and water molecules interacting with ATP (± Mg) are shown as sticks and coloured by heteroatom. Mg^2+^ coordination complex pseudo-bonds are show in purple dashed lines. See Figure S5B.
C) Unlike the monomeric or the dimer relay FICD mutants, dimeric wild-type FICD binds ATP in a configuration that would prevent BiP substrate AMPylation. The position of the α-phosphate in the FICD:ATP structure would preclude in-line nucleophilic attack (see Figure S5C-D). The left panel represents the superposition of the structures in the upper panel of *(B)*, with ATP interacting residues shown as sticks and annotated. Only Glu234 deviates significantly in sidechain position. Note, however, that the FICD:ATP His363 sidechain is also flipped, forming a hydrogen bond to a ribose interacting water (see *B*i). Mg^2+^ and ATP are coloured to match the corresponding ribbons. Active site waters are omitted for clarity. Distances are indicated by dashed black lines. The inset is a blow-up displaying distances *i-iv* between the γ-phosphates and Glu234 residues. Note, distances *i* and *ii* are derived from the γ-phosphate and Glu234 of different superimposed structures. Distances between Val316(Cγ1) and the corresponding Pα are shown in the right-hand side panel. See Figures S6-7.

**Table 1:** Data Collection and refinement statistics. Values in parentheses correspond to the highest-resolution shell, with the following exceptions: **\***The number of molecules in the biological unit is shown in parentheses; **\***\*MolProbity percentile score is shown in parentheses (100^th^ percentile is the best among structures of comparable resolutions, 0^th^ percentile is the worst).

|  | FICD:ATP | FICD<sup>K256S</sup>:Apo | FICD<sup>K256A</sup>:MgATP | FICD<sup>L258D</sup>:Apo | FICD<sup>L258D</sup>:MgATP | FICD<sup>L258D</sup>:MgAMP<br>-PNP |
| --- | --- | --- | --- | --- | --- | --- |
| **Data collection** |  |  |  |  |  |  |
| Synchrotron stations | DLS I04 | DLS I04 | DLS I03 | DLS I04 | DLS I03 | DLS I03 |
| Space group | *P*<sub>2</sub><sub>1</sub><sub>2</sub><sub>1</sub><sup>2</sup> | *P*<sub>22</sub><sub>1</sub><sub>2</sub><sub>1</sub> | *P*<sub>22</sub><sub>1</sub><sub>2</sub><sub>1</sub> | *P*<sub>3</sub><sub>1</sub><sub>2</sub><sub>1</sub> | *P*<sub>6</sub><sub>4</sub><sub>2</sub><sub>2</sub> | *P*<sub>6</sub><sub>4</sub><sub>2</sub><sub>2</sub> |
| Molecules in a.u.* | 2 (2) | 1 (2) | 1 (2) | 1 (1) | 1 (1) | 1 (1) |
| a,b,c; Å | 77.67, 107.65,<br>132.60 | 43.82, 76.51,<br>131.97 | 41.90, 73.98,<br>134.04 | 118.14, 118.14,<br>79.55 | 186.84, 186.84,<br>76.84 | 186.36, 186.36,<br>77.10 |
| $\alpha, \beta, \gamma; ^\circ$ | 90.00, 90.00, 90.00 | 90.00, 90.00, 90.00 | 90.00, 90.00, 90.00 | 90.00, 90.00, 120.00 | 90.00, 90.00, 120.00 | 90.00, 90.00, 120.00 |
| Resolution, Å | 83.58-2.70<br>(2.83-2.70) | 65.99-2.25<br>(2.32-2.25) | 134.04-2.32 (2.41-2.32) | 62.80-2.65 (2.72-2.65) | 93.42-2.54 (2.65-2.54) | 93.18-2.31 (2.39-2.31) |
| R<sub>merge</sub> | 0.163 (0.717) | 0.109 (0.385) | 0.107 (0.636) | 0.176 (0.856) | 0.167 (1.009) | 0.071 (0.611) |
| $\langle I/\sigma(I) \rangle$ | 19.2 (1.8) | 6.8 (2.4) | 5.6 (1.0) | 8.6 (2.2) | 13.0 (2.5) | 10.3 (1.8) |
| CC1/2 | 0.999 (0.720) | 0.993 (0.547) | 0.995 (0.567) | 0.996 (0.549) | 0.999 (0.503) | 0.998 (0.523) |
| No. of unique reflections | 31293 (4091) | 21825 (1978) | 18543 (1712) | 18963 (1380) | 26617 (3188) | 34573 (3351) |
| Completeness, % | 100.0 (100.0) | 99.9 (99.5) | 99.4 (97.3) | 100.0 (100.0) | 100.0 (100.0) | 99.4 (99.1) |
| Redundancy | 6.4 (6.5) | 4.4 (4.4) | 3.7 (3.7) | 9.7 (10.0) | 16.1 (16.5) | 4.6 (4.6) |
| **Refinement** |  |  |  |  |  |  |
| R<sub>work</sub>/R<sub>free</sub> | 0.280 / 0.319 | 0.208 / 0.259 | 0.282 / 0.325 | 0.228 / 0.283 | 0.232 / 0.252 | 0.214 / 0.251 |
| No. of atoms (non-H) | 5650 | 2851 | 2731 | 2951 | 2828 | 2940 |
| Average B-factors, Å<sup>2</sup> | 55.3 | 42.5 | 54.6 | 50.9 | 58.2 | 56.4 |
| RMS Bond lengths, Å | 0.002 | 0.003 | 0.003 | 0.003 | 0.002 | 0.003 |
| RMS Bond angles, ° | 1.142 | 1.180 | 0.763 | 1.222 | 1.127 | 1.170 |
| Ramachandran favoured region, % | 96.5 | 98.5 | 98.2 | 97.9 | 98.5 | 99.4 |
| Ramachandran outliers, % | 0 | 0 | 0 | 0 | 0 | 0 |
| MolProbity score** | 1.33 (100<sup>th</sup>) | 0.86 (100<sup>th</sup>) | 0.74 (100<sup>th</sup>) | 0.99 (100<sup>th</sup>) | 0.97 (100<sup>th</sup>) | 0.99 (100<sup>th</sup>) |
| PDB code | 6i7g | 6i7h | 6i7i | 6i7j | 6i7k | 6i7l |

When co-crystallised with MgATP or MgAMPPNP the resulting FICD structures contained clear densities for nucleotide (Figure 5B and S5B). The AMPylation-biased FICD mutants also contained discernible, octahedrally coordinated Mg^2+^ ions (Figure 5Bii-iii and S5B). As noted in other Fic AMPylases, this Mg^2+^ was coordinated by the α-and β-phosphates of ATP/AMPPNP and Asp367 of the Fic motif (Xiao *et al*, 2010; Khater & Mohanty, 2015b; Bunney *et al*, 2014). Interestingly, in the dimeric wild-type FICD:ATP structure, crystallised in the presence of MgATP, there was no density that could be attributed to Mg^2+^ (Figure 5Bi). The only possible candidate for Mg^2+^ in this structure was a water density, located between all three phosphates, that fell in the Fic motif’s anion-hole – a position incompatible with Mg^2+^ coordination (Zheng *et al*, 2017).

Alignment of the nucleotide-bound structures revealed that ATP or AMPPNP were bound very differently by the wild-type dimer and the AMPylation-biased monomeric or dimer relay FICD mutants (Figure 5C and S5C). Concordantly, the RMSD of ATP between the wild-type FICD and monomeric FICD^L258D^ was 2.17 Å (and 2.23 Å for FICD^K256A^’s ATP). As previously observed in other ATP-bound Fic proteins that possess an inhibitory glutamate, the nucleotide in FICD:ATP was in an AMPylation non-competent conformation (Engel *et al*, 2012; Goepfert *et al*, 2013) that is unable to coordinate Mg^2+^; an essential ion for FICD-mediated AMPylation (Ham *et al*, 2014). Moreover, the position of the ATP α-phosphate precludes in-line nucleophilic attack (by the hydroxyl group of BiP’s Thr518) due to the proximity of the flap residue Val316 (Figure 5C and S5D). Furthermore, an attacking nucleophile in-line with Pα-O3α would be at a considerable distance from the catalytic His363 (required to deprotonate Thr518’s hydroxyl group) (Figure 5Bi, 5C and S5D).

By contrast, in the active sites of FICD^K256A^ or FICD^L258D^ MgATP and MgAMPPNP assumed AMPylation-competent conformations: their α-phosphates were in the canonical position (Figure S5E), as defined by AMPylation-active Fic proteins lacking inhibitory glutamates (Xiao *et al*, 2010; Engel *et al*, 2012; Goepfert *et al*, 2013; Bunney *et al*, 2014). As a result, in-line nucleophilic attack into the α-β-phosphoanhydride bond of ATP would not be sterically hindered and the Nε2 of His363 would be well positioned for general base catalysis (Figure 5C and S5C-D).

The presence of ATP in both dimeric wild-type FICD and monomeric FICD^L258D^ (although in different binding modes) is consonant with the DSF data (Figure 4F and S4E). Apart from Glu234, the residues directly interacting with ATP are similarly positioned in all structures (maximum RMSD 0.83 Å). However, considerable variability is observed in Glu234, with an RMSD of 4.20 Å between monomeric and dimeric wild-type ATP structures, which may hint at the basis of monomerisation-induced AMPylation competency. In ATP-bound structures the inhibitory glutamate is displaced from the respective apo ground-state position, in which it forms an inhibitory salt-bridge with Arg374: R_2_ of the Fic motif (Figure S6A). However, the displacement of the Glu234 side chain observed in the FICD:ATP structure (from its position in FICD:Apo; PDB 4u0u) would be insufficient for AMPylation-competent binding of the γ-phosphate of an ATP/AMPPNP (see distances i and ii, Figure 5C and S5C). This steric clash is relieved by the side chain conformations observed in the AMPylation-competent structures (see iii and iv, Figure 5C and S5C).

The findings above suggest that the AMPylation-biased FICD mutants attain their ability to competently bind MgATP by increased flexibility at the top of the α_inh_ and by extension through increased Glu234 dynamism. It is notable that all the nucleotide triphosphate-bound FICDs crystallised with intact dimer interfaces (Figure S6A and B). Moreover, with the exception of direct hydrogen bonds to mutated Lys256 side chains, in all FICD crystals the putative dimer relay hydrogen-bond network was maintained (Figure S6A). It seems likely that much of the monomerisation-linked conformational flexibility that facilitates binding of MgATP in solution cannot be trapped crystallographically. Nonetheless, comparing B-factors across the nucleotide triphosphate-bound FICD structures is informative: despite similar crystal packing (Figure S6B) the average residue B-factors, both in the dimerisation interface and near Glu234, positively correlated with the AMPylation activities of the respective mutants (Figure S7).

### ATP is an allosteric modulator of FICD

Given the conspicuous difference in the ATP binding modes observed between AMPylation-competent FICD mutants and the AMPylation-incompetent wild-type dimeric FICD, we were intrigued by the possibility that ATP may modulate other aspects of FICD enzymology and regulation.

In order to explore the effects of nucleotide on the different pre-AMPylation complexes formed between either dimeric or monomeric FICD and its co-substrate, ATP-bound BiP, we utilised BioLayer Interferometry (BLI). Biotinylated, client-binding-impaired, ATPase-defective BiP^T229A-V461F^ was made nucleotide free (Apo) and immobilised on a streptavidin biosensor. Its interactions with catalytically inactive, dimeric FICD^H363A^ or catalytically inactive, monomeric FICD^L258D-H363A^ were measured in the presence and absence of nucleotides. The binding of both monomeric and dimeric FICD to immobilised BiP was greatly enhanced by the pre-saturation of BiP with ATP (Figure 6A and S8A). This is consistent with ATP-bound BiP as the substrate for FICD-mediated AMPylation (Preissler *et al*, 2015b). Moreover, the binding signal produced by immobilised, ATP-bound BiP interacting with monomeric FICD^L258D-H363A^:Apo was significantly stronger than that produced from the corresponding dimeric FICD^H363A^:Apo analyte (Figure 6A). In contrast, AMPylated BiP bound more tightly to dimeric FICD^H363A^ than to monomeric FICD^L258D-H363A^ (forming a pre-deAMPylation complex, Figure S2G). These findings align with the role of dimeric FICD in deAMPylation and the monomer in AMPylation.

**Figure 6.**
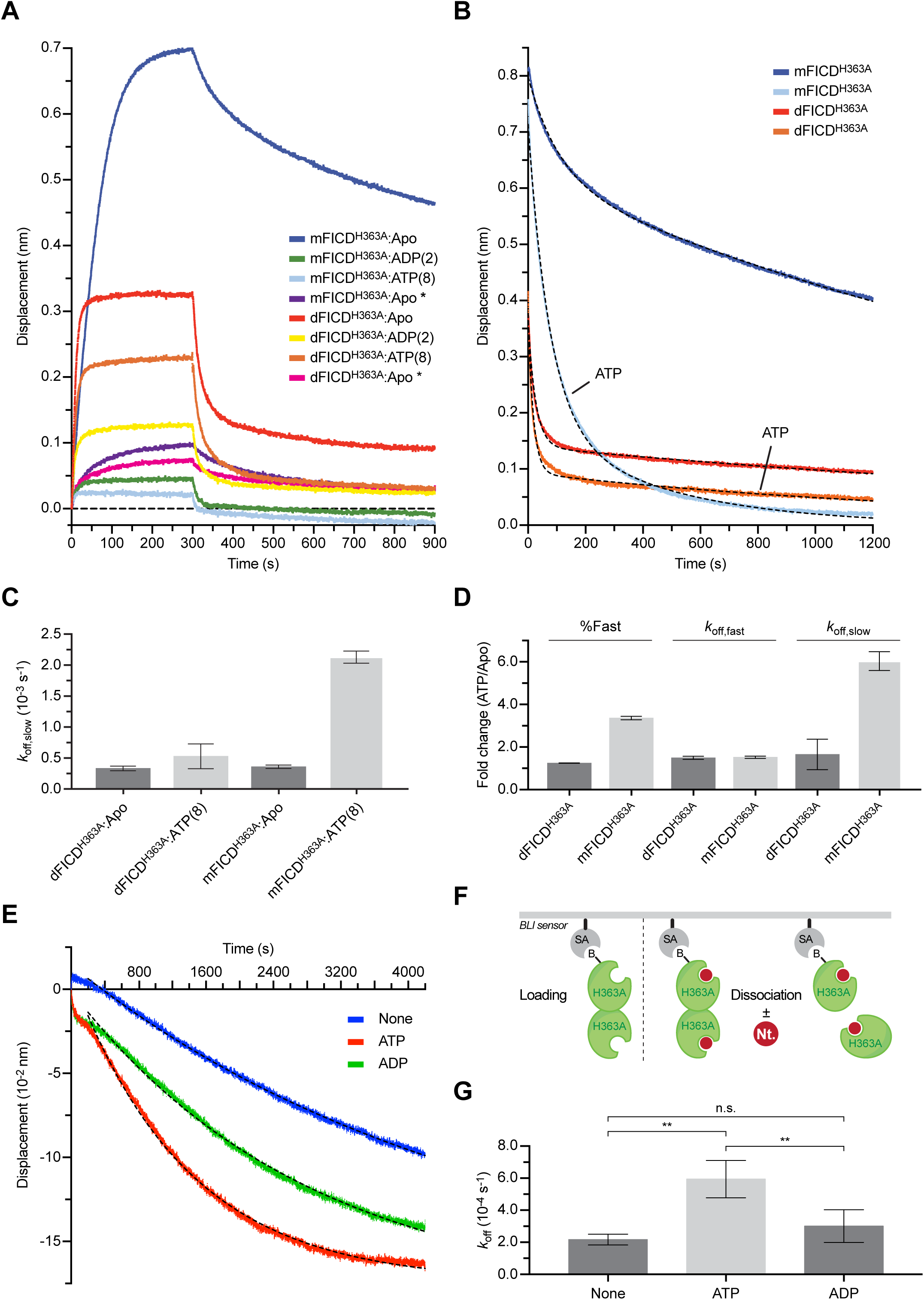
ATP destabilises the pre-AMPylation complex and the FICD dimer. A) BioLayer interferometry (BLI) derived association and dissociation traces of monomeric FICD^L258D-H363A^ (mFICD^H363A^) or dimeric FICD^H363A^ (dFICD^H363A^) from immobilized biotinylated BiP^T229A-V461F^ in absence or presence of nucleotides. Unless indicated (*) BiP was saturated with ATP before exposure to FICD variants. A representative experiment of three independent repetitions is shown. See Figure S8A-B.
B) BLI dissociation traces of proteins as in *(A)*. At t = 0 a pre-assembled complex of immobilised, ATP-saturated BiP and the indicated FICD proteins (associated without ATP) were transferred into a solution without or with ATP, as indicated. A representative experiment is shown and the biphasic dissociation kinetics are quantified in *(C)* and *(D)*. Full association and dissociation traces are shown in Figure S8C.
C) Graph of the slow dissociation rates (*k*_off,slow_) of monomeric FICD from BiP:ATP as shown in *(B)*. Bars represent mean values ± SD of three independent experiments.
D) The ATP-induced fold change in the percentage of the dissociation phase attributed to a fast dissociation (%Fast), *k*_off,fast_, and *k*_off,slow_ derived from the data represented in *(B)*. Bars show mean values ± SD of three independent experiments. See Figure S8D.
E) BLI dissociation traces of the FICD dimer at different nucleotide concentration. At t = 0 the species on the biosensor is a heterodimer of N-terminally biotinylated and an exchangeable, non-biotinylated FICD. Dissociation was conducted ± ligands (5 mM), as indicated. A representative experiment of four independent repeats, with mono-exponential fits are shown. See Figure S8E for raw data.
F) Cartoon schematic of the BLI assay workflow used to derive data presented in *(E)* and Figure S8E.
G) Quantification of the off rates derived from *(E)*. ATP, but no ADP, significantly increases the dimer dissociation rate [**: p < 0.01, by Tukey test; n.s.: not significant]. Data shown is the mean ± SD of four independent experiments.

Interestingly, in presence of magnesium bound nucleotide (either MgATP or MgADP) the FICD^H363A^ interaction with ATP-bound BiP was weakened (Figure 6A). This effect was considerably more pronounced for monomeric FICD^L258D-H363A^. To quantify the effect of FICD monomerisation on the kinetics of pre-AMPylation complex dissociation, BLI probes preassembled with biotinylated, ATP-bound BiP and either apo dimeric FICD^H363A^ or apo monomeric FICD^L258D-H363A^ were transferred into otherwise identical solutions ± ATP (schematised in Figure S8B). The ensuing dissociations fit biphasic exponential decays and revealed that ATP binding to FICD accelerated the dissociation of monomeric FICD^H363A^ more than dimeric FICD^H363A^ (Figure 6B and S8C). The effect of ATP was noted on both the slow dissociation phase of the monomer (*k*_off,slow_; Figure 6C-D) and on the percentage of dissociation attributed to the fast phase (%Fast; Figure 6D and S8D). The effect of ATP on the dissociation kinetics of the FICD^L258D-H363A^/BiP:ATP complex, measured under conditions of effectively infinite dilution, argues against a simple one-site competition between ATP-bound BiP and ATP for the Fic domain active site. Instead, these observations are better explained as allosteric modulation of monomeric FICD by ATP.

The structural data indicates that FICD’s oligomeric state can impact significantly on the mode of ATP binding, and Figure 6B indicates an allosteric effect of nucleotide binding on FICD. Together these observations suggested bi-directional intramolecular signalling from the dimer interface to the nucleotide-binding active site and therefore the possibility that ATP binding in FICD’s active site may also influence the oligomeric state of the protein. To investigate this hypothesis, hetero-dimers of N-terminally biotinylated FICD^H363A^ assembled with non-biotinylated FICD^H363A^ were loaded onto a BLI streptavidin biosensor. The dissociation of non-biotinylated FICD^H363A^ from its immobilised partner was then observed by infinite dilution into buffers varying in their nucleotide composition (Figure 6E and S8E, schematised in Figure 6F). ATP but not ADP induced a 3-fold increase in the dimer off rate (Figure 6G). This is suggestive of a mechanism whereby changing ATP/ADP ratios in the ER may modulate the oligomeric state of FICD.

## Discussion

This study addresses a key process in the post-translational UPR by which bifunctional FICD switches between catalysis of BiP AMPylation and deAMPylation, in order to match the folding capacity of the ER to the burden of unfolded proteins independently of changes in gene expression. The high affinity of FICD protomers for each other specifies the presence of principally dimeric FICD in the ER, shown here to restrict the enzyme to deAMPylation. This is the dominant mode of FICD both in vitro and in cells under basal conditions (Preissler *et al*, 2017a; Casey *et al*, 2017). However, establishing a pool of monomeric FICD unmasks its potential as a BiP AMPylase and enfeebles deAMPylation. The structural counterpart to this switch is the mode by which MgATP, the AMPylation reaction’s co-substrate, is productively engaged in the active site of the monomeric enzyme. Our studies suggest that monomerisation relieves the repression imposed on FICD AMPylation by weakening a network of intramolecular contacts. In the repressed state these contacts propagate from the dimer interface to the enzyme’s active site and stabilise a conserved inhibitory residue, Glu234, to block AMPylation-competent binding of MgATP (Figure 7).

**Figure 7.**
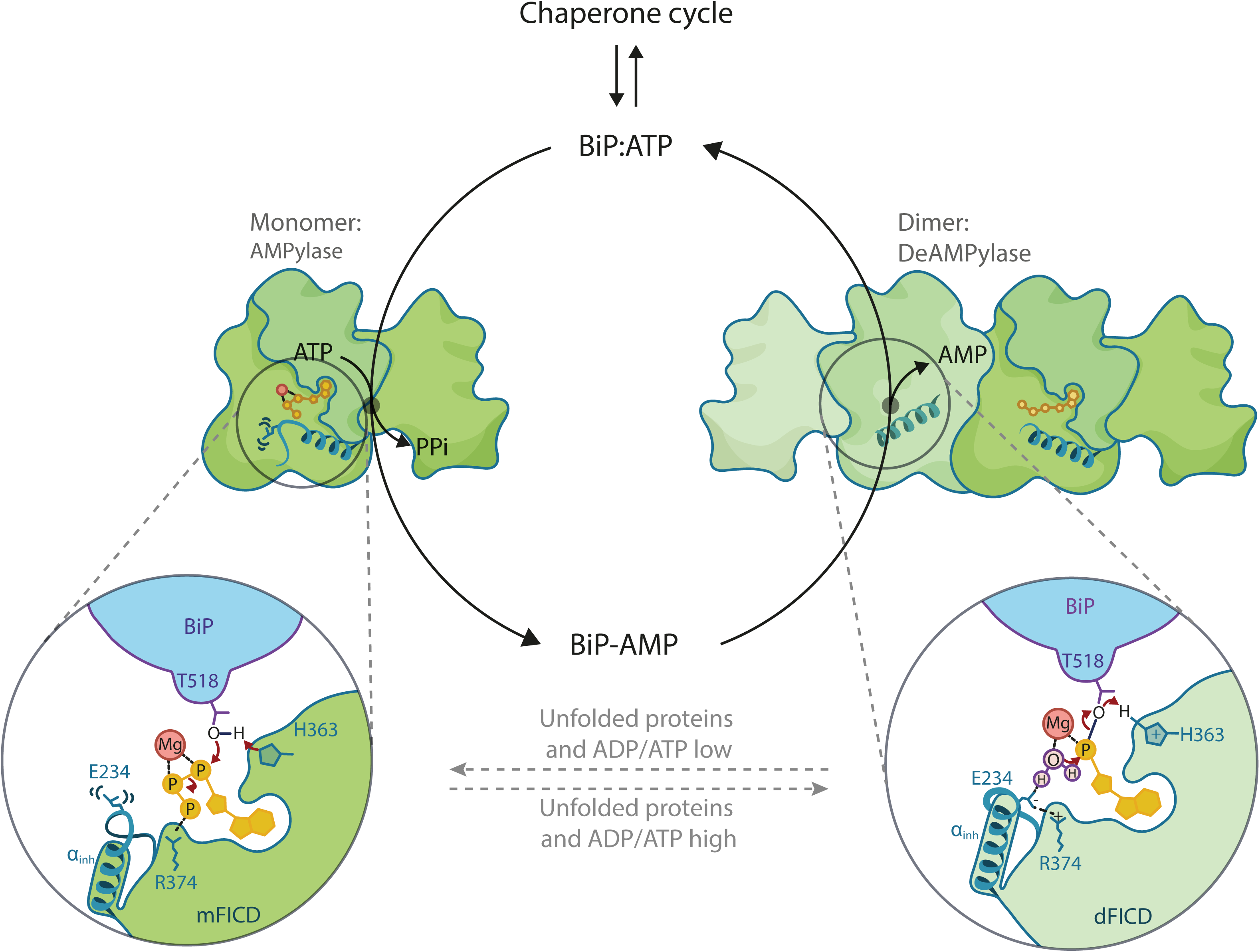
A proposed model of an oligomerisation state-dependent switch in FICD bifunctional active site. Under conditions of ER-stress the dimeric form FICD is favoured (right hand side). Dimeric FICD cannot bind ATP in an AMPylation competent mode but can efficiently catalyse deAMPylation of BiP-AMP (thereby remobilising BiP back into the chaperone cycle). A decrease in unfolded protein load in the ER, possibly associated with a decreased ER ADP/ATP ratio, shifts the FICD monomer-dimer equilibrium towards monomeric FICD. Monomeric FICD can bind MgATP in an AMPylation competent conformation and, as such, AMPylate and inactivate surplus BiP.

Our observations of a biphasic FICD concentration-dependent rescue of BiP AMPylation in *FICD*^-/-^ cells and the conspicuous ability of the monomerising Leu258Asp mutation to establish a modified BiP pool in *FICD*^-/-^ cells, all support an oligomeric state-dependent switch as a key contributor to FICD regulation in vivo. This case is further supported by the divergent enzymatic properties of monomeric mutants and enforced disulphide-linked dimers in vitro, and by measurements of the enzymatic activity of wild-type FICD in concentration regimes above and close to the dimerisation *K*_d_. Complete monomerisation resulted in a 19-fold increase in AMPylation activity and a 2-fold decrease in deAMPylation activity. The concordance between monomeric FICD^L258D^, dimerisation-defective mutants, and mutants in the repressive relay from the dimer interface to the active site gives confidence in the validity of the biophysical and structural insights provided by the mutants.

The inverse correlation observed between the thermal stability of FICD mutants and their AMPylation activity, supports a role for enhanced flexibility in enabling the enzyme to attain the conformation needed for catalysis of this reaction – a role clarified by the crystallographic findings (see below). The biophysical assays also suggest that monomeric FICD is more allosterically sensitive to ATP binding, as it exhibits a pronounced nucleotide-dependent reduction in the affinity for its co-substrate, ATP-bound BiP. The observation that ATP significantly accelerated the dissociation of monomeric, nucleotide-free FICD from ATP-bound BiP suggests that this feature of the monomer is mediated allosterically (not by enhanced susceptibility of a destabilised protein to co-substrate competition for the same active site). The lower affinity of monomeric FICD for its BiP:ATP co-substrate, in the context of a quaternary pre-AMPylation complex, conspicuously distinguishes it from the dimer and is a feature that may also enhance AMPylation rates: ground-state destabilisation has been demonstrated in a number of enzymes as a means of catalytic rate enhancement, by reducing the otherwise anti-catalytic tight binding of an enzyme to its substrate (Andrews *et al*, 2013; Ruben *et al*, 2013).

A structure of the quaternary pre-AMPylation complex, that could inform our understanding of the features of the monomeric enzyme, does not exist. Nevertheless, important insights into the effect of monomerisation were provided by structures of FICD and its nucleotide co-substrate. Dimeric wild-type FICD binds ATP (without magnesium) in an AMPylation incompetent mode. This is consistent with all other inhibitory glutamate containing Fic structures crystallised with ATP or ATP analogues (Engel *et al*, 2012; Goepfert *et al*, 2013). In stark contrast, we have discovered that despite the presence of an inhibitory glutamate, monomerisation, or mutations in residues linking the dimer interface to Glu234, permit the binding of ATP with magnesium in a conformation competent for AMPylation.

Our studies suggest that the disparity in FICD’s ATP binding modes stems from a monomerisation-induced increase in Glu234 flexibility (mediated by weakening of the dimer relay). This increase in flexibility is reflected in relatively subtle changes in the Glu234 side chain position, B-factor increases in the respective crystal structures, and markedly lower melting temperature of FICD^K256A^ and FICD^L258D^ relative to the wild-type dimer.

It seems likely that in solution monomerisation allows greater flexibility in this dimer relay network, facilitating motion and possibly unfolding at the top of the Glu234 containing α-helix (α_inh_). Such considerations could explain the comparatively small differences in the position of Glu234, but stark differences in nucleotide conformation, observed between the dimeric wild-type and monomeric or dimer relay mutant structures. That is to say, in solution the mutants exhibit sufficiently increased Glu234 dynamics to permit binding of MgATP in a catalytically competent mode. However, the crystallisation process quite possibly favours rearrangements, including α_inh_ refolding and crystallographic reconstitution of the dimer interface, and convergence towards a low energy state (the one stabilised in solution by dimerisation). This then outweighs the energetic penalty of the resulting (crystallographically-induced) electronically or sterically strained carboxylate-carboxylate (Glu234-Glu263) or glutamate-phosphate contacts (Figure 5C and S5C). Crystallisation may therefore facilitate the apparent convergence of mutant FICD Glu234 conformations towards that imposed in solution by the dimer. By contrast, dimeric wild-type FICD is never able to bind MgATP competently, either in solution or in crystallo, due to its unperturbed allosteric dimer relay and consequently inflexible Glu234.

Oligomerisation state-mediated regulation of AMPylation is not unique to FICD. Tetramerisation of bacterial NmFic antagonises auto-AMPylation and AMPylation of its substrate, DNA gyrase (Stanger *et al*, 2016). Though the surfaces involved in oligomerisation of this class III Fic protein are different from that of FICD, these two repressive mechanisms converge on the state of their α_inh_s. As such, divergent Fic proteins potentially exploit, for regulatory purposes, an intrinsic metastability of this structurally conserved inhibitory α-helix (Garcia-Pino *et al*, 2008). Interestingly, the more extensive dimerisation surface of FICD (which contains Leu258 and is situated at the boundary of the Fic domain core and the N-terminal Fic domain extension) also acts as a structurally conserved dimer interface in other class II bacterial Fic proteins: CdFic (Dedic et al, 2016) and *Bacteroides thetaiotaomicron* (BtFic; PDB: 3cuc), but not in the monomeric *Shewanella oneidensis* Fic (SoFic) protein (Goepfert et al, 2013). Moreover, a His57Ala mutation in dimeric CdFic (which is structurally equivalent to FICD^K256A^) causes increased solvent accessibility and auto-AMPylation of a region homologous to the loop linking FICD’s Glu242-helix and the α_inh_ (Dedic *et al*, 2016). Despite differences in detail, these findings suggest the conservation of a repressive relay from the dimer interface to the active site of dimeric Fic proteins.

Our biophysical observations also suggest a reciprocal allosteric signal propagated from FICD’s nucleotide binding site back to the dimer interface; enhanced dimer dissociation was induced by ATP but not ADP. Consequently, it is tempting to speculate that FICD’s oligomeric state and hence enzymatic activity might be regulated by the ADP/ATP ratio in the ER. Under basal conditions, low ADP concentrations allow ATP to bind both the monomeric and dimeric pools of FICD, shifting the equilibrium towards the monomer and favouring BiP AMPylation. Stress conditions may increase ADP concentration in the ER (perhaps by increased ER chaperone ATPase activity). This increase would be proportionally much greater than the concomitant decrease in [ATP] (in terms of respective fold changes in concentration). The increased [ADP] would therefore be able to effectively compete with ATP for the monomer-dimer FICD pools and thereby shift the equilibrium back towards the BiP de-AMPylating FICD dimer.

The regulation of BiP by FICD-mediated AMPylation and deAMPylation provides the UPR with a rapid post-translational strand for matching the activity of a key ER chaperone to its client load. The simple biochemical mechanism proposed here for the requisite switch in FICD’s antagonistic activities parallels the regulation of the UPR transducers, PERK and IRE1, whose catalytically-active conformation is strictly linked to dimerisation (Dey *et al*, 2007; Lee *et al*, 2008). A simple correlation emerges, whereby ER stress favours dimerisation of UPR effectors, activating PERK and IRE1 to regulate gene expression and the FICD deAMPylase to recruit BiP into the chaperone cycle (possibly through an increased ER ADP/ATP ratio). Resolution of ER stress favours the inactive monomeric state of PERK and IRE1 and, as suggested here, the AMPylation-competent monomeric FICD (Figure 7).

## Accession Numbers

The FICD crystal structures have been deposited in the PDB with the following accession codes: 6i7g (FICD:ATP), 6i7h (FICD^K256S^:Apo), 6i7i (FICD^K256A^:MgATP), 6i7j (FICD^L258D^:Apo), 6i7k (FICD^L258D^:MgATP), and 6i7l (FICD^L258D^:MgAMP-PNP).

## Acknowledgements

We thank the Huntington lab for access to the Octet machine, Claudia Flandoli for scientific illustrations and the CIMR flow cytometry core facility team (Reiner Schulte, Chiara Cossetti and Gabriela Grondys-Kotarba). This work was supported by Wellcome Trust Principal Research Fellowship to D.R. (Wellcome 200848/Z/16/Z), Medical Research Council PhD programme funding to L.A.P. (MR/K50127X/1), a Wellcome Trust Principal Research Fellowship to R.J.R. (Wellcome 082961/Z/07/Z), and a Wellcome Trust Strategic Award to the Cambridge Institute for Medical Research (Wellcome 100140).

## Author contributions

L.A.P. co-led and conceived the project, designed and conducted the biophysical experiments, analysed and interpreted the data, purified and crystallised proteins, collected, analysed and interpreted the X-ray diffraction data, and wrote the manuscript. C.R. designed, conducted and interpreted the in vivo experiments and contributed to revising the manuscript. Y.Y. supervised crystallisation efforts as well as the collection and processing of the X-ray diffraction data, contributed to analysis and interpretation of the structural data and to revising the manuscript. L.N. contributed to the in vivo experiments. S.H.M. conducted the AUC experiments and analysed the AUC data, and contributed to revising the manuscript. R.J.R. contributed to analysis and interpretation of the structural data and to revising the manuscript. D.R. conceived and oversaw the project, interpreted the data, and wrote the manuscript. S.P. co-led and conceived the project, designed and conducted the biochemical experiments, analysed and interpreted the data, and wrote the manuscript.

## Materials and Methods

### Plasmid construction

The plasmids used in this study have been described previously or were generated by standard molecular cloning procedures and are listed in Table S2.

### Cell lines

All cells were grown on tissue culture dishes or multi-well plates (Corning) at 37 °C and 5% CO_2_. CHO-K1 cells (ATCC CCL-61) were phenotypically validated as proline auxotrophs and their *Cricetulus griseus* origin was confirmed by genomic sequencing. *CHOP::GFP* and *XBP1s::Turquoise* reporters were introduced sequentially under G418 and puromycin selection to generate the previously-described derivative CHO-K1 S21 clone (Sekine *et al*, 2016). The cells were cultured in Nutrient mixture F-12 Ham (Sigma) supplemented with 10% (v/v) serum (FetalClone II; HyClone), 1 × Penicillin-Streptomycin (Sigma), and 2 mM L-glutamine (Sigma). The CHO-K1 *FICD*^-/-^ cell line used in this study was described previously (Preissler *et al*, 2015b). HEK293T cells (ATCC CRL-3216) were cultured in Dulbecco’s Modified Eagle’s Medium (Sigma) supplemented as described above. Cell lines were subjected to random testing for mycoplasma contamination using the MycoAlert Mycoplasma Detection Kit (Lonza).

Experiments were performed at cell densities of 60-90% confluence. Where indicated, cells were treated with cycloheximide (Sigma) at 100 µg/ml diluted with fresh, pre-warmed medium and then applied to the cells by medium exchange.

### Mammalian cell lysates

Cell lysis was performed as described in (Preissler *et al*, 2015a) with modifications. In brief, mammalian cells were cultured on 10 cm dishes and treated as indicated and/or transfected using Lipofectamine LTX with 5 µg plasmid DNA, and allowed to grow for 24 to 40 h. Before lysis, the dishes were placed on ice, washed with ice-cold PBS, and cells were detached in PBS containing 1 mM ethylenediaminetetraacetic acid (EDTA) using a cell scraper. The cells were sedimented for 5 min at 370 × *g* at 4 °C and lysed in HG lysis buffer [20 mM HEPES-KOH pH 7.4, 150 mM NaCl, 2 mM MgCl_2_, 10 mM D-glucose, 10% (v/v) glycerol, 1% (v/v) Triton X-100] containing protease inhibitors (2 mM phenylmethylsulphonyl fluoride (PMSF), 4 µg/ml pepstatin, 4 µg/ml leupeptin, 8 µg/ml aprotinin) with 100 U/ml hexokinase (from *Saccharomyces cerevisiae* Type F-300; Sigma) for 10 min on ice. The lysates were cleared for 10 min at 21,000 × *g* at 4 °C. Bio-Rad protein assay reagent (BioRad) was used to determine the protein concentrations of lysates. For analysis by SDS-PAGE, SDS sample buffer was added to the lysates and proteins were denatured by heating for 10 min at 70 °C before separation on 12.5% SDS polyacrylamide gels. To detect endogenous BiP by native-PAGE the lysate samples were loaded immediately on native gels (see below).

### Native polyacrylamide gel electrophoresis (native-PAGE)

Non-denaturing native-PAGE was performed as described (Preissler *et al*, 2015a). Briefly, Tris-glycine polyacrylamide gels (4.5% stacking gel and a 7.5% separation gel) were used to separate proteins from mammalian cell lysates to detect BiP monomers and oligomers. The separation was performed in running buffer (25 mM Tris, 192 mM glycine, pH ∼8.8) at 120 V for 2 h. Afterwards, the proteins were transferred to a polyvinylidene difluoride (PVDF) membrane in blotting buffer (48 mM Tris, 39 mM glycine; pH ∼9.2) supplemented with 0.04 (w/v) SDS for 16 h at 30 V for immunodetection. The membrane was washed for 20 minutes in blotting buffer (without SDS) supplemented with 20% (v/v) methanol before blocking. Volumes of lysates corresponding to 30 µg of total protein were loaded per lane to detect endogenous BiP from cell lysates by immunoblotting.

### Streptavidin pull-down and FLAG immunoprecipitation

To analyse the formation of FICD dimers in vivo (Figure 1B), CHO-K1 cells were transfected with 4 μg plasmid DNA encoding His_6_-AviTag-FICD (UK 2275) or His_6_-AviTag-FICD^L258D^ (UK 2319) and FLAG-FICD (UK 2276) or FLAG-FICD^L258D^ (UK 2318), and 4 μg plasmid DNA encoding BirA (in order to keep the final amount of plasmid DNA the same, an empty pCEFL plasmid was used; Table S2) as described above. 24 h before lysis the medium was exchanged to medium containing 50 μM

Biotin (Molecular Probes). For streptavidin pull-down of His_6_-AviTag-FICD, CHO-K1 cells were transfected and allowed to grow for approximately 40 h before lysis in lysis buffer [50 mM Tris-HCl pH 7.4, 150 mM NaCl, 1% (v/v) Triton X-100, 10% (v/v) glycerol] supplemented with protease inhibitors. The lysates were cleared twice, normalized and equal volumes of the lysates were incubated with 50 μl Dynabeads (MyOne Streptavidin C1, Life Technologies) for 60 to 90 min at 4 °C, rotating. The beads were then recovered by centrifugation for 1 min at 200 × *g* and by placing the tube in a magnetic separation stand. They were then washed three times at 25 °C with RIPA buffer [50 mM Tris-HCl pH 8.0, 150 mM NaCl, 1% (v/v) Triton X-100, 0.5% (v/v) sodium deoxycholate, 0.1% (v/v) SDS] supplemented with protease inhibitors. Bound proteins were eluted in 25 μl urea sample buffer [8 M urea, 1.36% (v/v) SDS, 12% (v/v) glycerol, 40 mM Tris-HCl pH 6.8, 0.002% (w/v) bromophenol blue, 100 mM DTT] and heating for 10 min at 70 °C. Equal volumes of the samples were loaded on a 12.5% SDS polyacrylamide gel and His_6_-AviTag-FICD and FLAG-FICD were detected by immunoblotting. Samples of the normalized lysates (60 μg) were loaded as an ‘input’ control.

For the reciprocal experiment, FLAG M2 immunoprecipitation of FLAG-FICD, equal volumes of the cleared and normalized lysates were incubated with 20 μl of Anti-FLAG M2 affinity gel (Sigma) for 60 to 90 min at 4 °C, rotating. The beads were then recovered by centrifugation for 1 min at 5,000 × *g* and washed three times with RIPA buffer. The proteins were eluted in 35 μl 2 × SDS sample buffer (without DTT) for 10 min at 70 °C. The beads were then sedimented and the supernatants were transferred to new tubes to which 50 mM DTT was added. Equal sample volumes were analysed by SDS-PAGE and immunoblotting as described above.

### Immunoblot analysis

After separation by SDS-PAGE or native-PAGE (see above) the proteins were transferred onto PVDF membranes. The membranes were blocked with 5% (w/v) dried skimmed milk in TBS (25 mM Tris-HCl pH 7.5, 150 mM NaCl) and incubated with primary antibodies followed by IRDye fluorescently labelled secondary antibodies (LI-COR). The membranes were scanned with an Odyssey near-infrared imager (LI-COR). Primary antibodies and antisera against hamster BiP [chicken anti-BiP (Avezov *et al*, 2013)], eIF2α [mouse anti-eIF2α (Scorsone *et al*, 1987)], FICD [chicken anti-FICD (Preissler *et al*, 2015b)], monoclonal anti-FLAG M2 (Sigma), and IRDye 800CW Streptavidin (LI-COR) were used.

### Flow cytometry

FICD (wild-type and mutants) over-expression-dependent induction of unfolded protein response signalling was analysed by transient transfection of wild-type and *FICD^-/-^ CHOP::GFP* CHO-K1 UPR reporter cell lines with plasmid DNA encoding the FICD protein and mCherry as a transfection marker, using Lipofectamine LTX as described previously (Preissler *et al*, 2015b). 0.5 µg DNA was used in Figure S2B-C to transfect cells growing in 12-well plates. 40 h after transfection the cells were washed with PBS and collected in PBS containing 4 mM EDTA, and single cell fluorescent signals (20,000/sample) were analysed by dual-channel flow cytometry with an LSRFortessa cell analyser (BD Biosciences). GFP and mCherry fluorescence was detected with excitation laser 488 nm, filter 530/30, and excitation laser 561, filter 610/20, respectively. Data were processed using FlowJo and median reporter (in Q1 and Q2) analysis was performed using Prism 6.0e (GraphPad).

### Production of VSV-G retrovirus in HEK293T cells and infection of CHO-K1 cells

In an attempt to establish BiP AMPylation in *FICD*^-/-^ cells (Figure 1A), cells were targeted with retrovirus expressing FICD (incorporating the naturally-occurring repressive uORF found in its cDNA) and mCherry. HEK293T cells were split onto 6 cm dishes 24 h prior to co-transfection of pBABE-mCherry plasmid encoding FICD (UK 1939; Table S2) with VSV-G retroviral packaging vectors, using TransIT-293 Transfection Reagent (Mirus) according to the manufacturer’s instructions. 16 h after transfection, medium was changed to medium supplemented with 1% (w/v) BSA (Sigma). Retroviral infections were performed following a 24 h incubation by diluting 0.45 µm filter-sterilized cell culture supernatants at a 1:1 ratio into CHO-K1 cell medium supplemented with 10 µg/ml polybrene (8 ml final volume) and adding this preparation to *FICD*^-/-^ CHO-K1 cells (1 × 10^6^ cells seeded onto 10 cm dishes 24 h prior to infection). Infections proceeded for 8 h, after which viral supernatant was replaced with fresh medium. 48 h later, the cells were split into four 10 cm dishes. Five days after transfection, single cells were sorted according to their mCherry intensity. Selected clones were expanded and analysed by flow cytometry (to assess mCherry intensity) and native-PAGE (to check for BiP AMPylation).

### Protein purification

#### FICD

Wild-type and mutant human FICD proteins (aa 104-445) were expressed as His_6_-Smt3 fusion constructs in T7 Express *lysY/I^q^* (NEB) *E. coli* cells. The cells were grown in LB medium (usually 6 l per construct) containing 50 µg/ml kanamycin at 37 °C to an optical density (OD_600nm_) of 0.6 and then shifted to 18 °C for 20 min, followed by induction of protein expression with 0.5 mM isopropylthio β-D-1-galactopyranoside (IPTG). The cultures were further incubated for 16 h at 18 °C, harvested, and lysed with a high-pressure homogenizer (EmulsiFlex-C3; Avestin) in buffer A [25 mM Tris-HCl pH 8.0, 500 mM NaCl, 40 mM imidazole, 1 mM MgCl_2_, 0.1 mM tris(2-carboxyethyl)phosphine (TCEP)] containing protease inhibitors [2 mM PMSF, 4 µg/ml pepstatin, 4 µg/ml leupeptin, 8 µg/ml aprotinin], 0.1 mg/ml DNaseI, and 20 µg/ml RNaseA. The lysates were centrifuged for 30 min at 45,000 × *g* and incubated with 1 ml of Ni-NTA agarose (Qiagen) per 1 l expression culture, for 30 min rotating at 4 °C. Afterwards, the beads were transferred to a gravity-flow Econo column (49 ml volume; BioRad), washed with five column volumes (CV) buffer A without MgCl_2_ and buffer B (25 mM Tris-HCl pH 8.0, 150 mM NaCl, 10 mM imidazole, 0.1 mM TCEP). The beads were further washed sequentially with buffer B sequentially supplemented with (i) 1 M NaCl, (ii) 10 mM MgCl_2_ + 5 mM ATP and (iii) 0.5 M Tris-HCl pH 8.0 [each 5 CV], followed by 2 CV TNT-Iz10 (25 mM Tris-HCl pH 8.0, 150 mM NaCl, 1 mM TCEP, 10 mM imidazole). Proteins were eluted by on-column cleavage with 1.5 µg/ml Ulp1 protease carrying a C-terminal StrepII-tag [Ulp1-StrepII (UK 1983)] in 1 bed volume TNT-Iz10 overnight at 4 °C. The eluate was collected, retained cleavage products were washed off the beads with TNT-Iz10, and all fractions were pooled. The total eluate was diluted 1:2 with 25 mM Tris-HCl pH 8.0 and further purified by anion exchange chromatography using a 6 ml RESOURCE Q column (GE Healthcare) equilibrated in 95% AEX-A (25 mM Tris-HCl pH 8.0, 25 mM NaCl) and 5% AEX-B (25 mM Tris-HCl, 1 M NaCl). Proteins were eluted by applying a gradient from 5-30% AEX-B in 20 CV at 3 ml/min. Fractions of elution peaks (absorbance at 280 nm, A_280nm_) corresponding to monomeric or dimeric FICD were pooled and concentrated using 30 kDa MWCO centrifugal filters (Amicon Ultra; Merck Millipore) in the presence of 1 mM TCEP. The proteins were then subjected to size-exclusion chromatography using a HiLoad 16/60 Superdex 200 prep grade column (GE Healthcare) equilibrated in SEC buffer (25 mM Tris-HCl pH 8.0, 150 mM NaCl). Peaks corresponding to monomeric or dimeric FICD were supplemented with 1 mM TCEP, concentrated (> 120 µM), and frozen in aliquots.

#### BiP

Mutant Chinese hamster BiP proteins with an N-terminal His_6_-tag were purified as described before with modifications (Preissler *et al*, 2017b). Proteins were expressed in M15 *Escherichia coli* (*E. coli*) cells (Qiagen). The bacterial cultures were grown in LB medium supplemented with 100 µg/ml ampicillin and 50 µg/ml kanamycin at 37 ° C to an OD_600nm_ of 0.8 and expression was induced with 1 mM IPTG. The cells were further grown for 6 h at 37 °C, harvested and lysed in buffer C [50 mM Tris-HCl pH 8, 500 mM NaCl, 1 mM MgCl_2_, 10% (v/v) glycerol, 20 mM imidazole] containing 0.1 mg/ml DNaseI and protease inhibitors. The lysates were cleared for 30 min at 45,000 × *g* and incubated with 1 ml of Ni-NTA agarose (Quiagen) per 1 l of expression culture, for 2 h rotating at 4 °C. Afterwards, the matrix was transferred to a gravity-flow Econo column (49 ml volume; BioRad) and washed with buffer D [50 mM Tris-HCl pH 8.0, 500 mM NaCl, 10% (v/v) glycerol, 30 mM imidazole], buffer E [50 mM Tris-HCl pH 8.0, 300 mM NaCl, 10 mM imidazole, 5 mM β-mercaptoethanol], and buffer E sequentially supplemented with (i) 1 M NaCl, (ii) 10 mM MgCl_2_ + 3 mM ATP, (iii) 0.5 M Tris-HCl pH 7.4, or (iv) 35 mM imidazole. The BiP proteins were then eluted with buffer F [50 mM Tris-HCl pH 8.0, 300 mM NaCl, 5 mM β-mercaptoethanol, 250 mM imidazole], dialyzed against HKM (50 mM HEPES-KOH pH 7.4, 150 mM KCl, 10 mM MgCl_2_) and concentrated with 30 kDa MWCO centrifugal filters. The proteins were flash-frozen in aliquots and stored at −80 °C.

GST-TEV-BiP constructs were purified like His_6_-Smt3-FICD, above, with minor alterations. Purification proceeded without the inclusion of imidazole in the purification buffers. Cleared lysates were supplemented with 1 mM DTT and incubated with GSH-Sepharose 4B matrix (GE Healthcare) for 1 h at 4 °C. 2 CV of TNT(0.1) (25 mM Tris-HCl pH 8.0, 150 mM NaCl, 0.1 mM TCEP) was used as a final wash step before elution. GST-TEV-BiP was eluted with 10 mM HEPES-KOH pH 7.4, 20 mM Tris-HCl pH 8.0, 30 mM KCl, 120 mM NaCl, 2 mM MgCl_2_ and 40 mM reduced glutathione. The eluate was cleaved with TEV protease (1/200 w/w; UK 759), whilst dialysing into TN plus 1 mM DTT, for 16 h at 4 °C. Uncleaved BiP was depleted by incubation, for 1 h at 4 °C, with GSH-Sepharose 4B matrix (1 ml per 5 mg of protein). The flow through was collected. Retained, cleaved material was washed from the matrix with 5 CV of TNT(0.1). All the cleaved, non-bound material was pooled. In order to AMPylate BiP, the cleaved product was combined with 1/50 (w/w) GST-TEV-FICD(45-458)^E234G^ (UK 1479; purified like the GST-TEV-BiP without TEV cleavage steps). The AMPylation reaction was supplemented with 10 mM MgATP (10 mM MgCl_2_ + 10 mM ATP), and incubated for 16 h at 25 °C. GST-TEV-FICD was then depleted by incubation with GSH-Sepharose 4B matrix, as above. Proteins were concentrated to > 200 µM. Aliquots were snap-frozen in liquid nitrogen and stored at −80 °C.

When required, protein samples were validated as being nucleotide free (Apo) by their A_260/280_ ratio and reference to IP-RP-HPLC analysis as conducted in (Preissler *et al*, 2017a).

#### Formation of disulphide-linked FICD dimers

Expression and purification of disulphide-linked dimers [of FICD^A252C-C421S^ (UK 2219) and FICD^A252C-H363A-C421S^ (trap; UK 2269)] was performed as described above with some alterations. After the affinity chromatography step, on-column cleavage was performed in TNT-Iz10 containing 1.5 µg/ml Upl1-StrepII and the retained cleavage products were washed off the beads with TN-Iz10 (25 mM Tris-HCl pH 8.0, 150 mM NaCl, 10 mM imidazole) in the absence of reducing agent. The pooled eluate was concentrated and diluted 1:4 with TN-Iz10. To allow for efficient disulphide bond formation the samples were supplemented with 20 mM oxidized glutathione and incubated overnight at 4 °C. Afterwards, the protein solutions were diluted 1:2 with 25 mM Tris-HCl pH 8.0 and further purified by anion-exchange and size-exclusion chromatography. The final preparations were analysed by non-reducing SDS-PAGE to confirm quantitative formation of covalently linked dimers (> 95%). Cysteine-free FICD^C421S^ (UK 2161) was purified according to the same protocol. A separate preparation of non-disulphide-bonded FICD^A252C-C421S^ (UK 2219), which was not subjected to oxidation with glutathione, was used in control experiments (Figure 3B, S3A and C).

### In vitro AMPylation

Standard radioactive in vitro AMPylation reactions were performed in HKMC buffer (50 mM HEPES-KOH pH 7.4, 150 mM KCl, 10 mM MgCl_2_, 1 mM CaCl_2_) containing 40 µM ATP, 0.034 MBq [α-^32^P]-ATP (EasyTide; Perkin Elmer), 0.2 µM FICD, and 1.5 µM ATP-hydrolysis and substrate-binding deficient BiP^T229A-V461F^ (UK 1825) in a final volume of 15 µl. Where indicated, samples contained 5 µM _S-S_FICD^A252C-H363A-C421S^ (UK 2269, trap) to sequester modified BiP. The reactions were started by addition of nucleotides. After a 20 min incubation at 25 °C the reactions were stopped by addition of 5 µl 4 × SDS sample buffer and denaturation for 5 min at 75 °C. The samples were applied to SDS-PAGE and the gels were stained with Coomassie (InstantBlue; expedeon). The dried gels were exposed to a storage phosphor screen and radioactive signals were detected with a Typhoon biomolecular imager (GE Healthcare). Signals were quantified using ImageJ64 software (NIH).

The reactions to analyse AMPylation at elevated concentrations (2 or 10 µM; Figure S4A) contained 2 µM BiP^T229A-V461F^, 80 µM ATP and 0.034 MBq [α-^32^P]-ATP in a final volume of 15 µl. The reactions were stopped after 5 min incubation at 25 °C.

Time course experiments (Figure 2E) were performed likewise but reactions contained 40 µM ATP, 0.136 MBq [α-^32^P]-ATP, 0.3 µM FICD, 2 µM BiP^T229A-V461F^, and 5 µM trap in a final volume of 60 µl. The reactions were incubated at 30 °C and samples (15 µl) were taken at different time intervals and processed as described above.

To study the effect of the concentration of wild-type FICD protein on its ability to establish a pool of AMPylated BiP (Figure 3A), final reactions were setup with 400 µM ATP, 0.049 MBq [α-^32^P]-ATP, 2.5 nM to 400 nM FICD (UK 2052) and 5 µM BiP^T229A-V461F^, without or with 5 µM trap in a final volume of 15 µl. The reactions were pre-incubated for 2 h before addition of nucleotides. After 2 and 16 h incubation with nucleotides at 25 °C (as indicated) samples (5 µl) were taken and denatured by heating in SDS sample buffer for analysis.

To compare the activity of disulphide-bonded FICD under non-reducing and reducing conditions (Figure 3C) _S-S_FICD^A252C-C421S^ protein (UK 2219) was pre-incubated 16 h at 25 °C without or with 10 mM DTT and a sample was analysed by non-reducing SDS-PAGE after denaturation in SDS sample buffer containing 40 mM N-ethylmaleimide (NEM). Afterwards, AMPylation reactions (15 µl final volume) were set up with 400 µM ATP, 0.049 MBq [α-^32^P]-ATP, 2.5 nM to 400 nM _S-S_FICD^A252C-C421S^, and 5 µM BiP^T229A-V461F^ in the presence or absence of 5 mM DTT. Samples were incubated for 16 h at 25 °C and 5 µl was taken and processed for analysis by reducing SDS-PAGE as described above. Parallel reactions performed with cysteine-free FICD^C421S^ (UK 2161), which underwent the same purification and oxidation procedure, served as a control. The experiment presented in Figure S3C was performed accordingly under non-reducing conditions, but the reactions were incubated for 2 h at 25 °C and in the presence of 5 µM trap.

### Coupled in vitro AMPylation/deAMPylation reactions

To measure AMPylation-/deAMPylation-dependent AMP production by FICD proteins reactions were set up in HKM buffer containing 250 µM ATP, 0.0185 MBq [α-^32^P]-ATP, 3 mM TCEP, 5 µM ATP-hydrolysis-deficient BiP^T229A^ (UK 838), and 2 µM FICD proteins in a final volume of 30 µl. The reactions were started by addition of nucleotides and incubated for 2 h at 30 °C. Afterwards, 2 µl were spotted onto a thin layer chromatography (TLC) plate (PEI Cellulose F; Merck Millipore) pre-spotted with 2 µl of nucleotide mix containing AMP, ADP, and ATP (each at 3.5 mM). The TLC plate was developed with 400 mM LiCl and 10% (v/v) acetic acid as a mobile phase and the dried plates were exposed to a storage phosphor screen. The signals were detected with a Typhoon biomolecular imager and quantified using ImageJ64.

### DeAMPylation measured by fluorescence polarisation (FP)

Measurement of deAMPylation kinetics was performed as described previously (Preissler *et al*, 2017a) with modifications. The probe (BiP^V461F^ modified with fluorescent, FAM-labelled AMP; BiP^V461F^-AMP^FAM^) was generated by pre-incubating FICD^E234G^ at 25 µM in HKM buffer with 200 µM ATP-FAM [N^6^-(6-amino)hexyl-adenosine-5’-triphosphate; Jena Bioscience] for 10 min at 30 °C, followed by addition of 25 µM His_6_-tagged BiP^V461F^ (UK 182) to a final volume of 50 µl, and further incubation for 2 h at 30 °C. Afterwards, the reaction was diluted with 950 µl of HKMG-Iz20 [50 mM HEPES-KOH pH 7.4, 150 mM KCl, 10 mM MgCl_2_, 5% (v/v) glycerol, 20 mM imidazole] and BiP proteins were bound to 80 µl Ni-NTA agarose beads (Qiagen) for 30 min at 25 °C in the presence of 0.01% Triton X-100. Following several wash steps in the same buffer proteins were eluted in HKMG-Iz250 [50 mM HEPES-KOH pH 7.4, 150 mM KCl, 10 mM MgCl_2_, 5% (v/v) glycerol, 250 mM imidazole], flash-frozen in aliquots, and stored at −80 °C.

DeAMPylation reactions were performed in FP buffer [50 mM HEPES-KOH pH 7.4, 150 mM KCl, 10 mM MgCl_2_, 1 mM CaCl_2_, 0.1% (v/v) Triton X-100] in 384-well polysterene microplates (black, flat bottom, µCLEAR; greiner bio-one) at 30 °C in a final volume of 30 µl containing trace amounts of fluorescent BiP^V461F-AMP-FAM^ probe (17 nM) and FICD proteins (0.75 or 7.5 µM). Fluorescence polarisation of FAM (λ_ex_ = 485 nm, λ_em_ = 535 nm) was measured with an Infinite F500 plate reader (Tecan). Fitting of the raw data to a single-exponential decay function was done using Prism 6.0e (GraphPad).

### Analytical size-exclusion chromatography

Analytical size-exclusion chromatography (SEC) was performed as described previously (Preissler *et al*, 2015a). Purified FICD proteins were adjusted to 20 µM in HKMC buffer (50 mM HEPES-KOH pH 7.4, 150 mM KCl, 10 mM MgCl_2_, 1 mM CaCl_2_) and incubated at 25 °C for at least 20 min before injection. From each sample 10 µl was injected onto a SEC-3 HPLC column (300 Å pore size; Agilent Technologies) equilibrated with HKMC at a flow rate of 0.3 ml/min. Runs were performed at 25 °C and A_280nm_ absorbance traces were recorded. Protein standards (Bio-Rad, cat. no. 151– 1901) were run as size references and the elution peaks of γ-globulin (158 kDa), ovalbumin (44 kDa), and myoglobulin (17 kDa) are indicated. For dimer SEC studies in Figure S1D-E, the FICD proteins were incubated for 16 h at 25 °C before injection. To investigate capture of AMPylated BiP by _S-S_FICD^A252C-H363A-C421S^ (UK 2269, trap), by SEC (Figure S2H), in vitro AMPylation reactions containing different combinations of 20 µM BiP^T229A-V461F^ (UK 1825), 10 µM trap, and 3 µM FICD^L258D^ (UK 2091) were performed in HKMC (supplemented with 2 mM ATP when indicated) and incubated for 1.5 h at 30 °C before injection.

### Fluorescence detection system sedimentation velocity analytical ultracentrifugation (FDS-SV-AUC)

Bacterial expression and purification of FICD proteins carrying an N-terminal cysteine for site-specific labelling (FICD^NC^, UK 2339, and FICD^L258D-NC^, UK 2367) was performed as described above with the following alterations: Cells were lysed in the presence of 5 mM β-mercaptoethanol and the eluate pool after affinity chromatography and on-column cleavage was supplemented with 5 mM DTT and diluted 1:2 with 25 mM Tris-HCl pH 8.0 containing 0.2 mM TCEP. The subsequent anion-exchange chromatography step was performed with buffer solutions AEX-A and AEX-B supplemented with 0.2 mM TCEP. Afterwards, the peak fractions corresponding to the dimeric form of FICD were pooled and concentrated. The protein at 200 µM was labelled in 150 µl with 600 µM Oregon Green 488-iodoacetamide in the presence of 0.5 mM TCEP and 0.1 mM EDTA for 16 h at 4 °C. The reaction was quenched with 2 mM DTT for 10 min at 25 °C. Afterwards the sample was passed through a CentriPure P2 desalting column (emp) equilibrated in SEC buffer containing 0.2 mM TCEP. The eluate was applied to size-exclusion chromatography using a Superdex 200 10/300 GL column (GE Healthcare) in the presence of 0.2 mM TCEP. The fractions of the A_280nm_ peak, corresponding to dimeric FICD, were pooled and the concentration of TCEP was adjusted to 1 mM. The proteins were concentrated and frozen in aliquots. The protein concentration was determined after denaturing the proteins with 6 M guanidine hydrochloride by measuring absorbance at 280 nm and 496 nm with a NanoDrop Spectrophotometer (Thermo Fisher Scientific). The concentration was calculated using the following equation:

Protein concentration (M) = [A_280nm_ – (A_496nm_ × 0.12)]/ε

Where 0.12 is the correction factor for the fluorophore’s absorbance at 280 nm, and ε is the calculated molar extinction coefficient of FICD (29,340 cm^-1^M^-1^). The labelling efficiency of the FICD^NC^ preparation was 74% as calculated based on the A_496nm_ value and assuming an extinction coefficient for Oregon Green 488 of 70,000 cm^-1^M^-1^. The labelling efficiency of the monomeric FICD^L258D-NC^ control preparation was 9.6%. Labelling of the endogenous cysteine residue (Cys421) of wild-type FICD was very inefficient (< 1%) and thus considered negligible.

Samples of Oregon Green-labelled FICD in 50 mM HEPES-KOH pH 7.4, 150 mM KCl, 10 mM MgCl_2_, 1 mM CaCl_2_, 0.3 mM TCEP, 0.1 % Tween-20, 0.15 mg/ml BSA (Sigma), ranging in concentration from 1.6 µM to 31 pM, were centrifuged at 45,0000 rpm at 20 °C in an An50Ti rotor using an Optima XL-I analytical ultracentrifuge (Beckmann) equipped with a fluorescence optical detection system (Aviv Biomedical) with fixed excitation at 488 nm and fluorescence detection at > 505 nm. Data were processed and analysed using SEDFIT 15 and SEDPHAT 13b (Schuck, 2003) according to the published protocol for high-affinity interactions detected by fluorescence (Chaturvedi *et al*, 2017). Data were plotted with Prism 6.0e (GraphPad) or GUSSI (Brautigam, 2015).

### Differential Scanning Fluorimetry (DSF)

DSF experiments were performed on an ABi 7500 qPCR machine (Applied Biosciences). Experiments were carried out in 96-well qPCR plates (Thermofisher), with each sample in technical triplicate and in a final volume of 20 µl. Protein was used at a final concentration of 2 µM, ligands at the concentration indicated in the figure legend (2.5-20 mM), and SYPRO Orange (Thermofisher) dye at a 10x concentration in a buffer of HKM plus 1 mM TCEP (unless otherwise specified). For the ATP titration (Figure 4F and S4E), the DSF buffer was supplemented with an additional 15 mM MgCl_2_ (25 mM total MgCl_2_). Fluorescence of the SYPRO Orange dye was monitored over a temperature range of 20-95 °C using the VIC filter set. Data was then analysed in Prism 7.0e (GraphPad), with melting temperature calculated as the global minimums of the negative first derivatives of the relative fluorescent unit melt curves (with respect to temperature).

### Bio-layer interferometry (BLI)

#### In vitro biotinylation

Ligands for BLI were generated from the tag cleaved forms of unmodified or AMPylated GST-TEV-AviTag-haBiPV461F(19-654) (UK 2043) and GST-TEV-AviTag-haBiP(28-635)^T229A-V461F^ (UK 2331). Biotinylation was conducted in vitro on 100 µM target protein, with 200 µM biotin (Sigma), 2 µM GST-BirA (UK 1801) in a buffer of 2 mM ATP, 5 mM MgCl_2_, 25 mM Tris-HCl pH 8.0, 150 mM NaCl and 1 mM TCEP. The reaction mixture was incubated for 16 h at 4 °C. Excess biotin was removed by size-exclusion chromatography on a S200 10/300 GL column (GE Healthcare) with a distal 1 ml GSTrap 4B (GE Healthcare), connected in series. The ligand was confirmed as being > 95% biotinylated as judged by streptavidin gel-shift. In the case of Biotinylated-AviTag-haBiP(28-635)^T229A-V461F^ this protein was also made nucleotide free by the addition of 2 U CIP (NEB) per mg of BiP, plus extensive dialysis into TN buffer with 1 mM DTT and 2 mM EDTA (dialysed with several dialysate changes, for 2 days at 4 °C). The protein was then purified by anion exchange chromatography on a MonoQ 5/50 GL column (GE Healthcare) using buffers AEX-A and AEX-B with a gradient of 7.5-50% B over 20 CV at a flow rate of 1 ml/min. The protein was concentrated using a 30 kDa MWCO centrifugal filters (Amicon Ultra; Merck Millipore) and then gel filtered, as above, but into an HKM buffer. Fractions were pooled and supplemented with 1 mM TCEP. All proteins after biotinylation and purification were concentrated to > 20 µM, flash-frozen in small aliquots and stored at −80 °C.

#### Kinetic experiments

All BLI experiments were conducted on the FortéBio Octet RED96 System (Pall FortéBio) in a buffer basis of HKM plus 0.05% Triton X-100 (HKMTx). Nucleotide was added as indicated. Streptavidin (SA)-coated biosensors (Pall FortéBio) were hydrated in HKMTx for at least 30 min prior to use. All BLI experiments were conducted at 30 °C with the experimental steps as indicated in the text. BLI reactions were prepared in 200 µl volumes in 96 well microplates (greiner bio-one, cat. no. 655209). Ligand loading was performed for 300 to 600 s at a shake speed of 1000 rpm until a binding signal of 1 nm was reached. The immobilised ligand sensor was then baselined in assay solution for at least 200 s. For kinetic experiments with biotinylated-AviTag-haBiP(28-635)^T229A-V461F^:Apo [BiP^T229A-V461F^:Apo (UK 2331)] loaded on the tip, a 10 Hz acquisition rate was used and the baseline, association and dissociation steps were conducted at a 400 rpm shake speed. Preceding the baseline step biotinylated BiP^T229A-V461F^:Apo was also activated with or without ATP (2 mM unless otherwise stated), as indicated, for 300 s at a 1000 rpm shake speed. In these experiments FICD analyte association or dissociation steps were conducted in the presence or absence of nucleotide, as indicated, with ATP at 8 mM and ADP at 2 mM. These concentrations were chosen in an attempt to saturate either monomeric or dimeric FICD with the respective nucleotide [*K*_d_ of MgADP for wild-type FICD is 1.52 µM by ITC (Bunney *et al*, 2014); *K*_1/2_ of ATP induced FICD T_m_ shift in the low mM range] and/or to make ATP binding non-rate limiting. In Figure S8A, as a control for the absence of substantial ATP dissociation from BiP, between the activation and baseline step an additional 1500 s BiP wash (± ATP) was included, as indicated. Other BLI experiments were conducted with all steps at a 1000 rpm shake speed with a 5 Hz acquisition rate. All association-dissociation kinetics were completed in ≤ 1500 s. Data was processed in Prism 7.0e (GraphPad). Note, the FICD variants used as analytes in all BLI experiments were catalytically inactive His363Ala variants (used at 250 nM).

In the dimer dissociation BLI experiments biotinylated AviTag-FICD(104-458)^H363A^ (UK 2422) was diluted to 3 nM and incubated for 10 min at 25 °C with either dimeric FICD^H363A^ or monomeric FICD^L258D-H363A^ (at 300 nM) in HKMTx. After this incubation period the streptavidin biosensors were loaded until those immobilising hetero-labelled dimers (biotinylated AviTag-FICD(104-458)^H363A^ with FICD^H363A^) were loaded to a 1 nm displacement. Dissociation was initiated by dipping in HKTx buffer (50 mM HEPES-KOH pH 7.4, 150 mM KCl and 0.05% Triton X-100) ± nucleotide at 5 mM, as indicated. Data was processed by subtracting the respective monomer incubated biotinylated FICD tip from the dimeric hetero-labelled dimer dissociation, followed by fitting of the corrected dissociation to mono-exponential decay using Prism 7.0e (GraphPad).

### Protein crystallization and structure determination

FICD proteins were purified as above in *Protein Purification* but gel filtered into a final buffer of 10 mM Tris-HCl pH 8.0, 150 mM NaCl and 1 mM TCEP [T(10)NT]. Proteins were diluted to 9 mg/ml in T(10)NT prior to crystallisation, via sitting drop vapour diffusion. For structures containing ATP, final diluted protein solutions were supplemented with MgATP (from a pH 7.4, 100 mM stock solution) to a final concentration of 10 mM. A drop ratio of protein solution to crystallisation well solution of 200:100 nl was used. Where applicable crystals were obtained by microseeding (D’Arcy *et al*, 2007), from conditions provided in Table S1. In these instances, a drop ratio of protein solution to water-diluted seeds to crystallisation well solution of 150:50:100 nl was used. The best diffracting crystals were obtained in crystallisation conditions detailed in Table S1.

Diffraction data were collected from the Diamond Light Source, and the data processed using XDS (Kabsch, 2010) and the CCP4 module Aimless (Winn *et al*, 2011; Evans & Murshudov, 2013). Structures were solved by molecular replacement using the CCP4 module Phaser (McCoy *et al*, 2007; Winn *et al*, 2011). For the FICD^L258D^:Apo and FICD:ATP structures the human FICD protein (FICD:MgADP) structure 4U0U from the Protein Data Bank (PDB) was used as a search model. Subsequent molecular replacements used the solved FICD^L258D^:Apo structure as a search model. Manual model building was carried out in COOT (Emsley *et al*, 2010) and refined using refmac5 (Winn *et al*, 2003). Metal binding sites were validated using the CheckMyMetal server (Zheng *et al*, 2017). Polder (OMIT) maps were generated by using the Polder Map module of Phenix (Liebschner *et al*, 2017; Adams *et al*, 2010). Structural figures were prepared using UCSF Chimera (Pettersen *et al*, 2004) and PyMol (Schrödinger, LLC, 2015).

---

**Figure S1.**
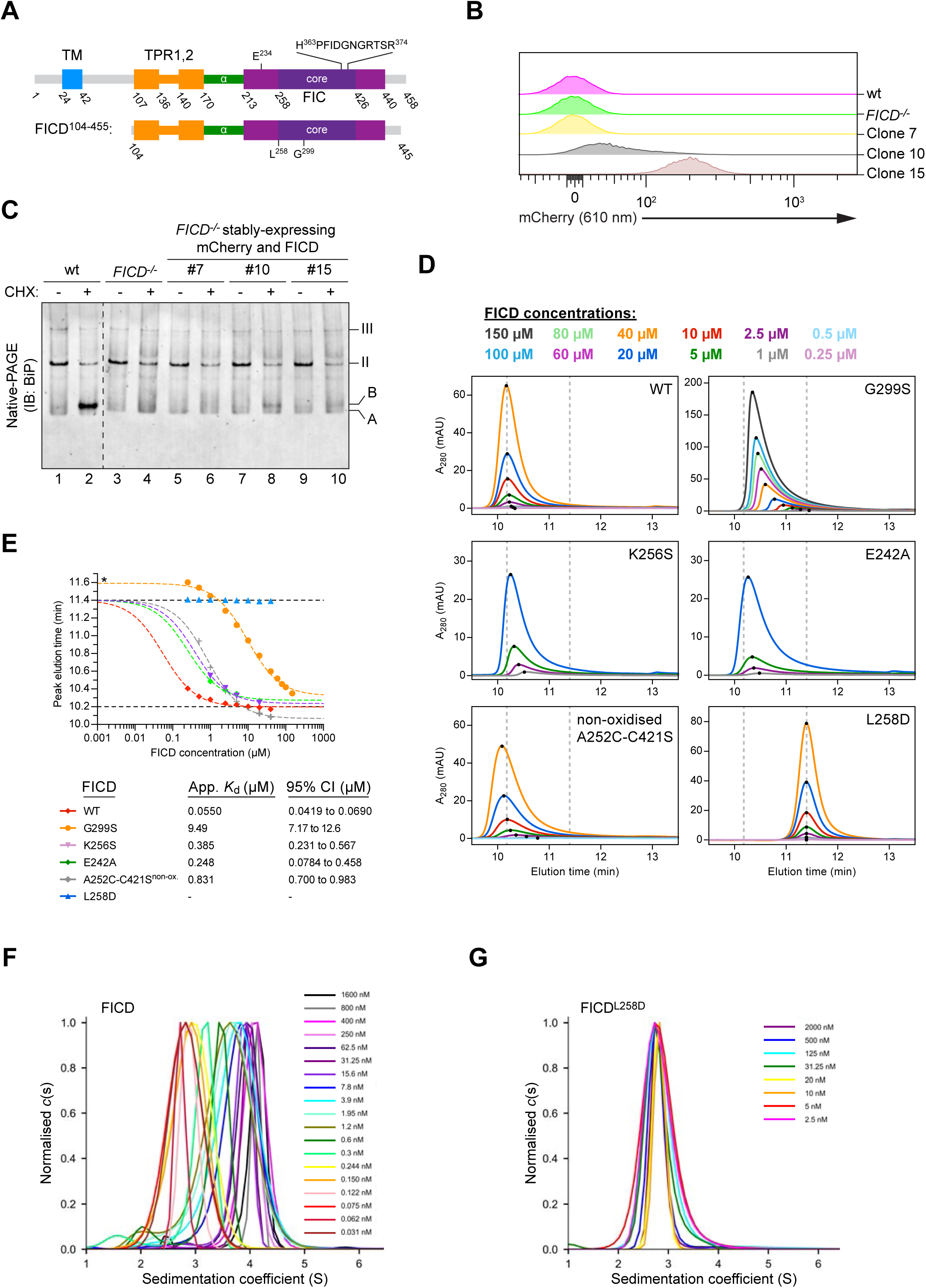
Low-level expression facilitates AMPylation in vivo and FICD mutations are able to disrupt the tight dimer formed in solution. A) Schematic representation of the domain organization of FICD and the shorter protein fragment used for in vitro experiments. The transmembrane domain (blue), the TPR domain (orange), the α-helical linker (green), the Fic domain (purple) and the core Fic domain (deep purple) including the active site motif are indicated.
B-C) Characterization of CHO-K1 *FICD*^-/-^ UPR reporter clones stably expressing wild-type FICD**. (B)** Flow cytometry analysis of CHO-K1 *FICD*^-/-^ UPR reporter clones stably-expressing mCherry and FICD. Clones were selected based on mCherry signal, assuming a direct correlation with FICD expression levels. **(C)** Immunoblot of endogenous BiP from CHO-K1 *FICD*^-/-^ clones shown in *(B)* exposed to cycloheximide as in Figure 1A. Note that only clone 10, with an intermediate mCherry signal, showed detectable accumulation of AMPylated BiP.
D-E) Size-exclusion chromatography (SEC) analysis of wild-type and mutant FICD proteins. **(D)** SEC elution profiles with FICD proteins at the indicated concentrations. Black dots mark the position of the elution peaks. Dotted lines mark the approximate elution peak times for dimeric (10.2 min) and monomeric (11.4 min) FICD, respectively. **(E)** Plot of the elution peak times from *(D)* as a function of protein concentration. With the exception of FICD^G299S^ (*; a mutation that shifts the elution time relative to the monomer) best-fit monomer-dimer association curves are shown with the top plateau constrained to the monomer elution time (11.4 min). Approximate dimerisation *K*_d_s were derived and are shown in the figure key for the different partially monomerising mutants (with 95% confidence intervals). Note that FICD^L258D^ eluted as a monomer and wild-type FICD principally as a dimer at all concentrations tested (0.2-50 µM). Conversely, FICD^G299S^ and non-oxidized FICD^A252C-C421S^ formed much weaker dimers. As in *(D)* the monomer and dimer elution times are represented by dotted (horizontal) lines.
F-G) Analysis of FICD by analytical ultracentrifugation. Overlays of *c*(*s*) distributions of **(F)** wild-type FICD and **(G)** FICD^L258D^ are shown in units of experimental *s*-values. A signal-weighted isotherm for the wild-type protein (Figure 1E) was generated from integration of the titration series distributions.

**Figure S2.**
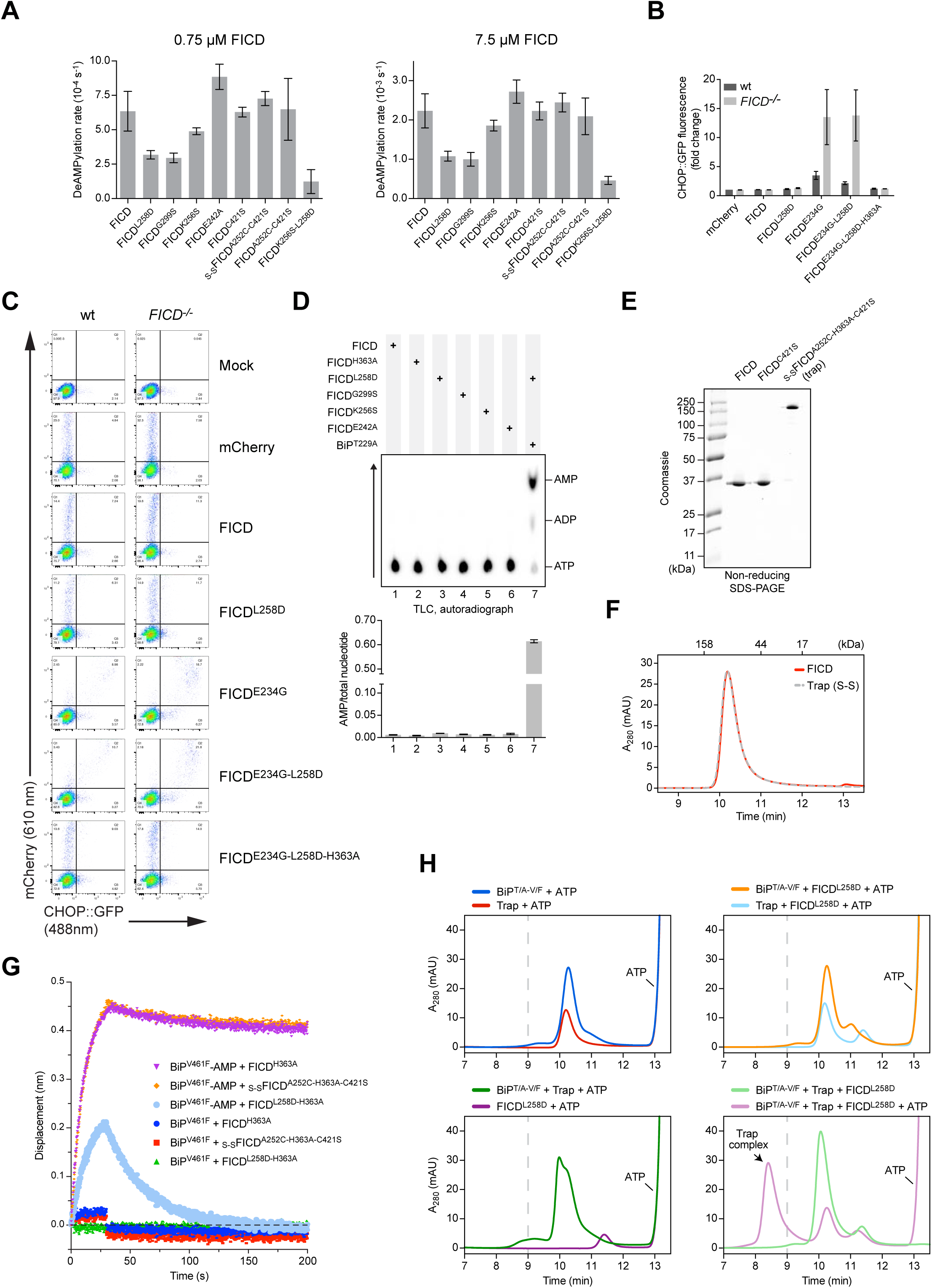
Monomerisation inhibits deAMPylation and markedly stimulates FICD AMPylation activity. A) Summary of deAMPylation rates of wild-type and mutant FICD proteins. Shown are deAMPylation rates of BiP^V461F^-AMP^FAM^ by the indicated FICD proteins (at 0.75 µM or 7.5 µM) as detected by a change in fluorescence polarisation. Mean values ± SD of the normalized raw data fitted to a single-exponential decay function of at least four independent measurements are presented.
B-C) The effect of FICD overexpression on a UPR reporter. **(B)** Flow cytometry analysis of wild-type and *FICD*^-/-^ CHO-K1 *CHOP::GFP* UPR reporter cells transfected with plasmids encoding wild-type or the indicated FICD derivatives and a mCherry transfection marker. Shown are the median values ± SD of the GFP fluorescence signal of mCherry-positive cells from three independent experiments (fold change relative to wild-type cells transfected with a plasmid encoding mCherry alone). Note that only Glu234Gly-containing, deAMPylation-deficient FICDs activate the reporter. **(C)** Flow cytometry raw data of a representative experiment quantified in *(B)*.
D) AMP production by FICD dimer interface or relay mutants is BiP dependent. AMP production in the presence of [α-^32^P]-ATP was measured by TLC and autoradiography (as in Figure 2B). Plotted below are mean AMP values ± SD (n = 3).
E-G) Characterization of covalently linked _S-S_FICD^A252C-H363A-C421S^ dimers – a trap for BiP-AMP. **(E)** Coomassie-stained, SDS-PAGE gel of the indicated FICD proteins. **(F)** Size-exclusion chromatography elution profiles of wild-type FICD and covalently linked _S-S_FICD^A252C-H363A-C421S^ (trap) dimers at 20 µM, as in Figure 1D. Note that the oxidised trap elutes, like the wild-type FICD, as a dimer. **(G)** BioLayer interferometry (BLI) derived association and dissociation traces of the indicated FICD proteins (in solution) from immobilized AMPylated (BiP-AMP) or unmodified BiP. The trap (_s-s_FICD^A252C-H363A-C421S^) and FICD^H363A^ had indistinguishable tight interaction with BiP-AMP (with low off rates). The interaction of BiP-AMP with monomeric FICD^L258D-H363A^ was more transient. The interaction between these FICD variants and unmodified BiP was further diminished.
H) Sequestration of AMPylated BiP by trap FICD analysed by SEC. Elution profiles of in vitro AMPylation reactions containing the indicated components in the presence • r absence of covalently linked _S-S_FICD^A252C-H363A-C421S^ (trap) dimers to sequester the AMPylated BiP product. Note that the trap forms a stable complex with BiP when AMPylated by monomeric FICD^L258D^. An early eluting species, representing a stable complex between modified BiP and trap, only occurs in the reaction containing AMPylation-active, monomeric FICD^L258D^ and ATP (bottom right panel, pink trace). Here, BiP-mediated ATP hydrolysis and substrate interactions were discouraged by use of a BiP^T229A-V461F^ double mutant.

**Figure S3.**
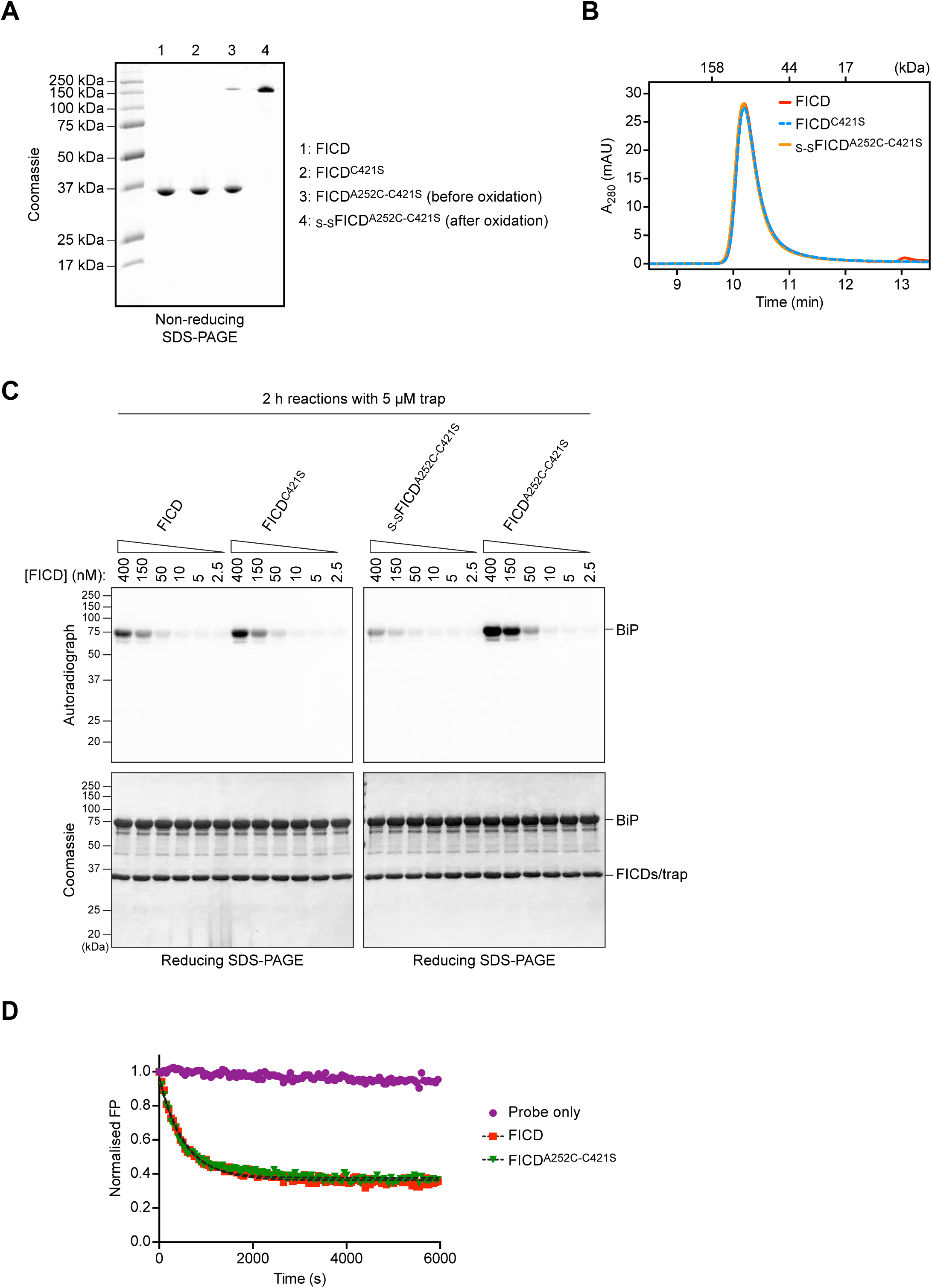
Non-disulphide-linked FICD^A252C-C421S^ shows enhanced AMPylation activity. A) Coomassie-stained, non-reducing SDS-PAGE gel of the indicated FICD proteins.
B) Size exclusion chromatography (SEC) elution profiles of FICD proteins injected at a concentration of 20 µM. Protein absorbance at 280 nm is plotted against elution time. The elution times of protein standards are indicated as a reference. Note that wild-type FICD, FICD^C421S^, and oxidised _S-S_FICD^A252C-C421S^ co-elute as dimers. See Figure S1D-E.
C) Radioactive in vitro AMPylation reactions were performed as in the right hand side panel of Figure 3A, that is with the indicated FICD proteins under non-reducing conditions in presence of covalently linked _S-S_FICD^A252C-H363A-C421S^ dimers (trap). Note that the accumulation of modified BiP correlates with the FICD concentration. Less modified BiP was produced by covalently-linked, oxidised _S-S_FICD^A252C-C421S^ dimers, whereas more AMPylated BiP was generated in reactions containing non-oxidised FICD^A252C-C421S^. The trap, present at 5 µM, co-migrates with the indicated FICD enzyme and dominates the signal in the Coomassie stained gel (FICD/trap).
D) Time-dependent in vitro deAMPylation of fluorescent BiP^V461F^-AMP^FAM^ by the indicated FICD proteins (at 7.5 µM) assayed by fluorescence polarisation (as in Figure 2A). A representative experiment (data points and fit curves) is shown and deAMPylation rates are presented in Figure S2A. Note that non-oxidised FICD^A252C-C421S^ has very similar deAMPylation kinetics to the wild-type protein. This contrasts with the oxidised form which displays a slight increase in deAMPylation rate (Figure 3D and S2A).

**Figure S4.**
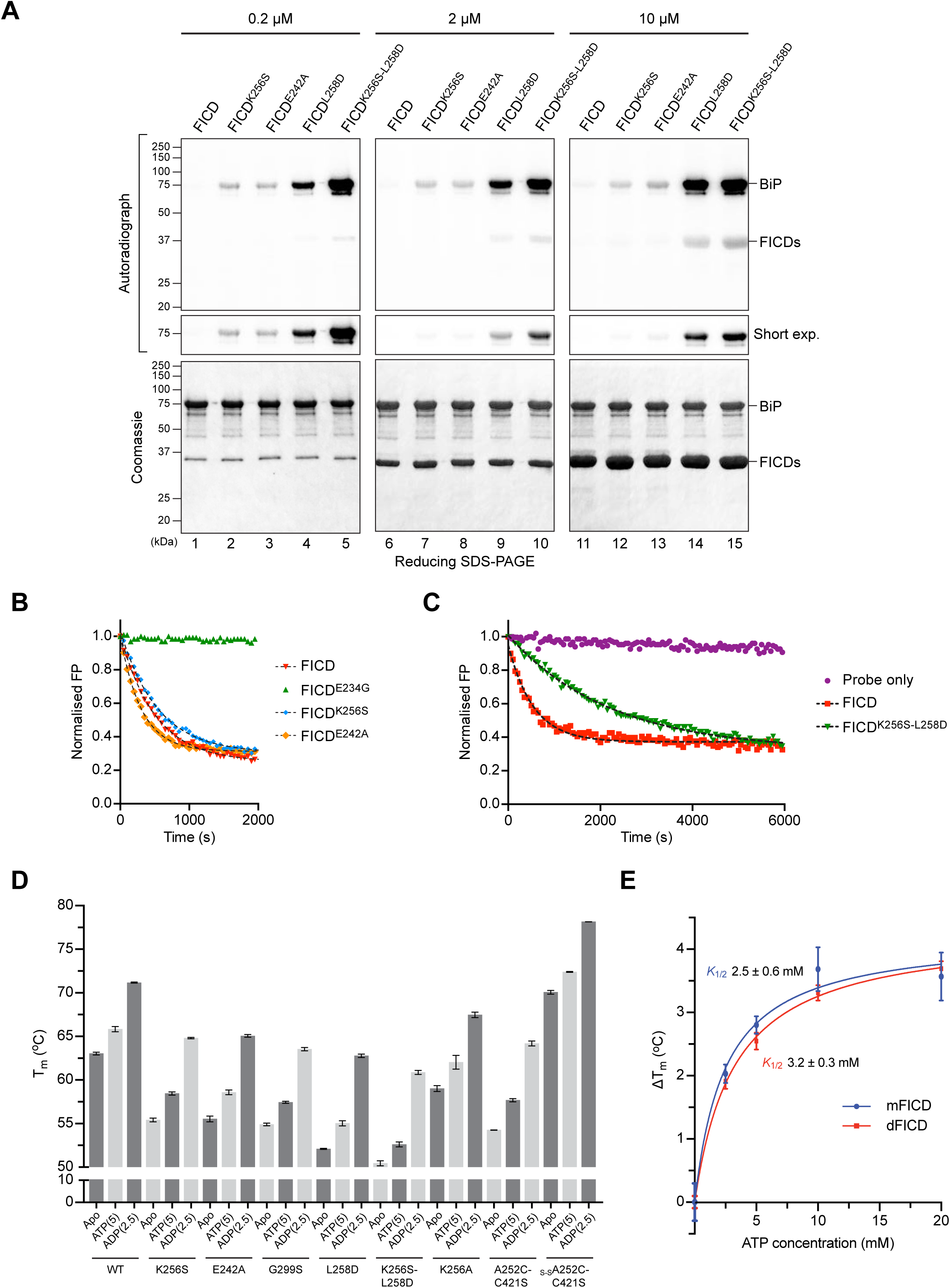
FICD dimer relay mutants produce a pool of AMPylated BiP in vitro, and FICD AMPylation activity correlates with increased flexibility. A) Radioactive in vitro AMPylation reactions with the indicated FICD proteins at the indicated concentrations, [α-^32^P]-ATP, and BiP^T229A-V461F^ were analysed by SDS-PAGE. The radioactive signals were detected by autoradiography and proteins were visualised by Coomassie staining. Note the enhanced production of AMPylated BiP in the presence of dimer relay mutants, FICD^K256S^ and FICD^E242A^, relative to the wild-type protein and a further increase in the production of AMPylated BiP by the monomeric FICD^K256S-L256D^ double mutant relative to the monomeric FICD^L258D^. Also note the auto-AMPylation signals of the monomeric FICDs at high enzyme concentration.
B-C) In vitro deAMPylation of fluorescent BiP^V461F^-AMP^FAM^ by the indicated FICD proteins (at 7.5 µM) measured by fluorescence polarisation. A representative experiment (data points and fit curves) is shown and rates are presented in Figure S2A. Note the impaired deAMPylation activity of the monomeric FICD^K256S-L256D^ double mutant in *(C)*.
D) DSF T_m_ analysis of wild-type (wt) and mutant FICD proteins in absence (Apo) or presence of ATP or ADP. Nucleotide concentrations are given in parentheses. Non-oxidised and oxidised forms of FICD^A252C-C421S^ were assayed in buffer lacking reducing agent (which did not affect the T_m_ of wild-type FICD; not shown). Shown are the mean T_m_ values ± SD from three independent experiments. Note that FICD^K256A^ is more stable than FICD^K256S^ but less than wild-type FICD. Furthermore, the stabilities of oxidised and non-oxidised FICD^C421S-A252C^ relative to the wild-type correlate inversely with their AMPylation activities (Figure 3B). The same data for the wild-type FICD, FICD^E242A^, FICD^G299S^, FICD^L258D^ and FICD^K256S-L258D^ in the Apo state are presented in Figure 4E.
E) Plot of the increase in FICD melting temperature (ΔT_m_) against ATP concentration as measured by DSF (derived from Figure 4F). Note the similarity in the plot of FICD^L258D^ (mFICD) and the wild-type dimer (dFICD); mFICD *K*_½_ 2.5 ± 0.6 mM and dFICD *K*_½_ 3.2 ± 0.3 mM. Shown are mean ΔT_m_ values ± SD of three independent experiments with the best fit lines for a one site binding model.

**Figure S5.**
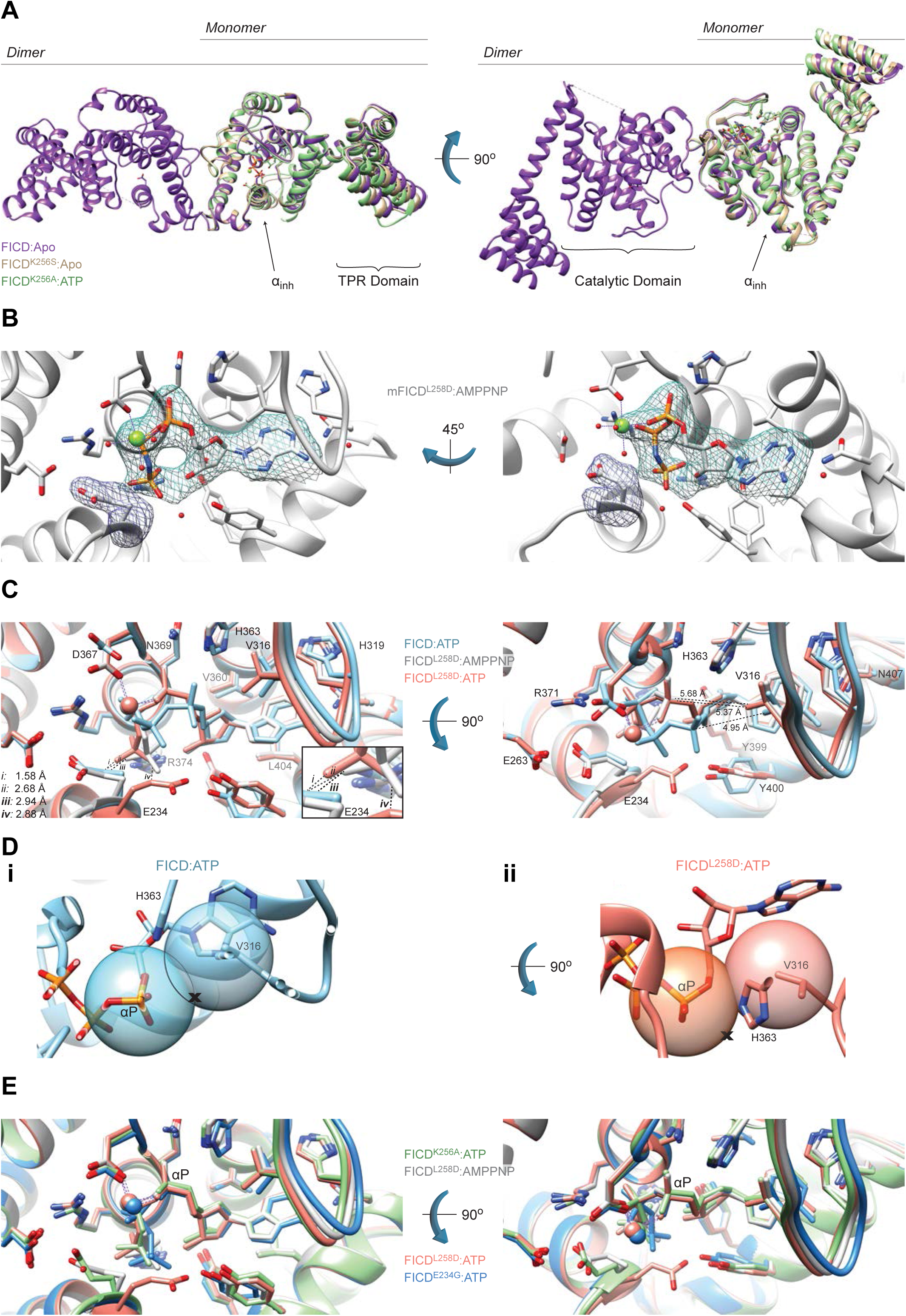
Monomerisation allows ATP to bind to FICD in a mode conducive to BiP AMPylation. A) Mutation of the dimer relay residue Lys256 does not result in large conformational changes in FICD. Shown is the alignment (residues 213-407) of the molecules in the asymmetric unit. Structures are coloured as indicated. Glu234, ATP and Mg (where applicable), are shown as sticks. The inhibitory alpha helix (α_inh_) and gross domain architecture is annotated. The FICD:Apo structure is from PDB: 4U0U.
B) Electron density of both MgAMPPNP and the inhibitory Glu234, from monomeric FICD^L258D^ co-crystallized with MgAMPPNP. Unbiased polder (OMIT) maps are shown in blue and purple meshes, contoured at 3.0 and 5.0 σ, respectively. All residues and water molecules interacting with MgAMPPNP are shown as sticks and coloured by heteroatom. Mg^2+^ coordination complex pseudo-bonds are show in purple dashed lines.
C) Unlike wild-type FICD, monomeric FICD^L258D^ binds ATP and ATP analogues in an AMPylation competent conformation. The indicated structures and distances are shown as in Figure 5C, with ATP interacting residues shown as sticks and annotated. The position of the α-phosphate relative to Val316 in the FICD:ATP structure (see distances in right hand side panel) would preclude in-line nucleophilic attack (see D-E). The inset is a blow-up displaying distances *i-iv* between the γ-phosphates and Glu234 residues. A potentially significant difference in the Glu234 position between the FICD^L258D^:MgAMPPNP and FICD:ATP structures is apparent: hypothetical distance *ii* (2.68 Å, between Glu234 of FICD:ATP and AMPPNP γ-phosphate of FICD^L258D^) is less favourable than the observed distance *iii* (2.94 Å, between the AMPPNP γ-phosphate and Glu234 of FICD^L258D^). Note, His363 of FICD:ATP is in a non-optimal flip state to facilitate general base catalysis (see Figure 5B).
D) *(i)* The mode of ATP binding in wild-type dimeric FICD sterically occludes the nucleophilic attack required for AMPylation. Shown are semi-opaque 3 Å centroids centred on Pα and Val316 (Cγ1). The putative BiP Thr518 nucleophile (depicted by the cross) is positioned in-line with the scissile phosphoanhydride (parallel to the plane of the paper) and 3 Å from Pα. This nucleophile position lies within the Val316 centroid (indicating a steric clash). For clarity, the FICD:ATP structure is overlaid with a thin slice of the FICD:ATP structure in the plane of the Pα-O3α bond. ***(ii)*** In the monomeric AMPylation-competent FICD^L258D^:ATP structure the nucleophile lies outside the Val316 centroid in proximity to His363 (the general base).
E) The ATP α-phosphates of monomer or dimer relay mutants are in the same position as that competently bound to the AMPylation unrestrained dimeric FICD^E234G^. Shown are all AMPylation competent MgATP structures overlaid as in *(C)* and Figure 5C. The dimeric FICD^E234G^:MgATP (dark blue, PDB: 4U07) is also included as a reference for an active AMPylating enzyme.

**Figure S6.**
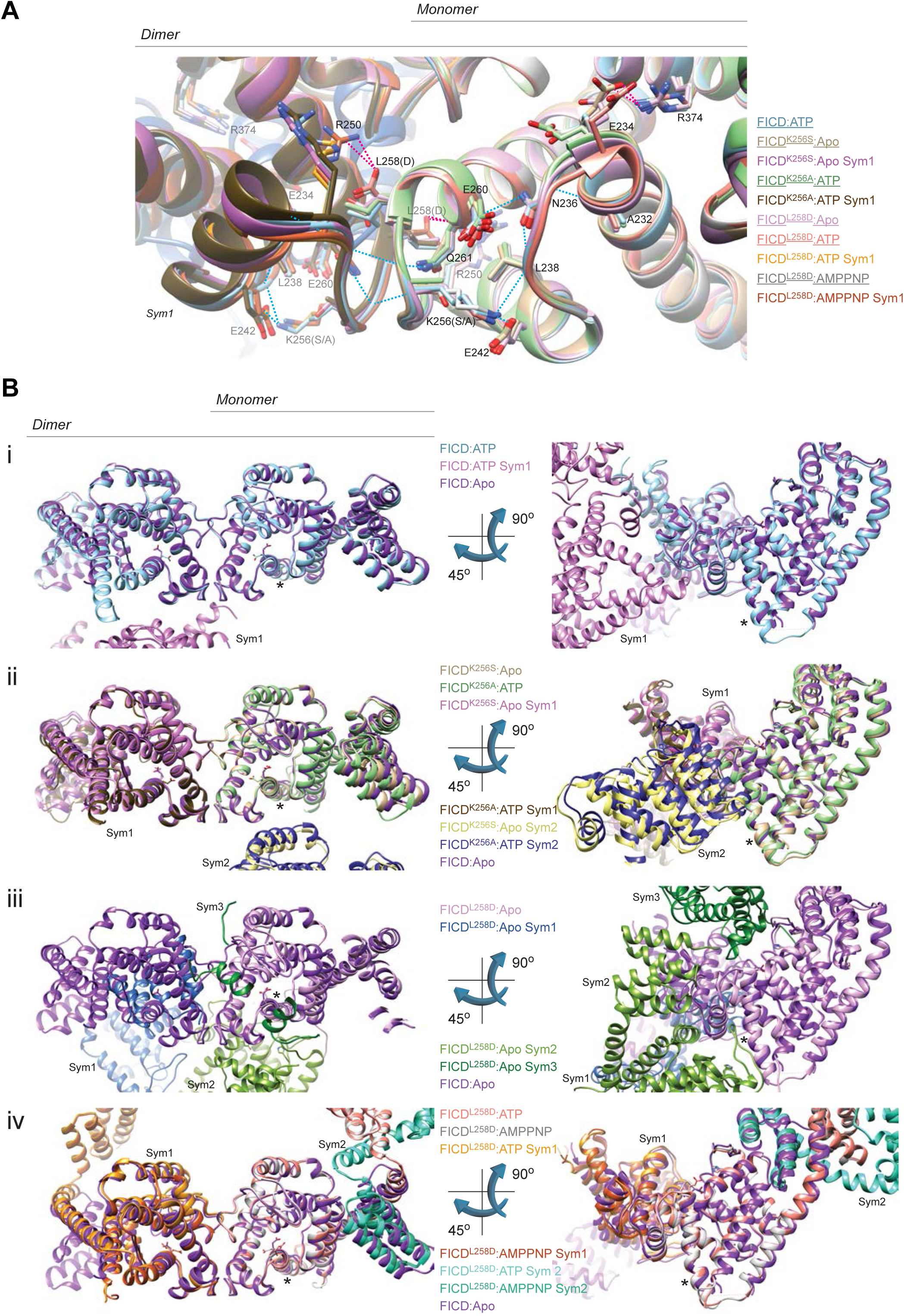
FICD crystallographic packing and dimer interface. A) The hydrogen bonding network connecting the dimer interface and enzyme active site is maintained in the crystal structures of monomeric and Lys256 mutant FICDs. An alignment of the hydrogen bond network linking the dimer interface to Glu234 in the indicated structures is displayed (in the same view as Figure 4A). H-bonds are shown in blue dashed lines. Where indicated, single molecules from the asymmetric unit (underlined) are displayed with their respective symmetry mates (Sym1). Note, the side-chains of Asp258 and (Sym1)Arg250 of the monomeric FICD^L258D^ (cocrystallised with nucleotide) form a crystallographically induced inter-molecular H-bond (magenta dashed line). The salt-bridges between the Glu234 and the Fic motif Arg374 (magenta dashed lines) in the FICD^L258D^:Apo and FICD^K256S^:Apo structures, observed in other inhibitory glutamate-containing Fic crystal structures, are also shown.
B) Dimer interface contacts are imposed crystallographically, and crystal packing around the a_inh_ is similar in all FICD structures. FICDs with similar crystal packings are grouped into panels *(i-iv)*. The inhibitory alpha helix (α_inh_) is denoted with an asterisk (*) and Glu234s are shown as sticks. The wild-type dimeric FICD:Apo structure (FICD:Apo; PDB:4U0U) is provided in all panels for reference. Where a single FICD molecule constituted the asymmetric unit, symmetry mates within 4 Å of its dimer interface (Sym1) or 4 Å of its inhibitory helix region (Sym2/3) are also displayed. Note that crystals of the Lys256 mutants *(ii)* contain a single molecule in their asymmetric unit but are packed as dimers, crystallographically reconstituting the dimeric biological unit. The asymmetric unit of FICD^L258D^ bound to ATP (or an ATP analogue) *(iv)* contains a single molecule and thus corresponds to the biological unit of this monomeric protein. However, packing against its symmetry mates (Sym1), crystallographically reconstitutes a dimer interface that is highly similar, but not identical, to that observed in the wild-type protein (see Asp258 and (Sym1)Arg250 in S6A, above). Sym2 in *(iv)* serves to highlight the replacement of the flipped out TPR domain with the flipped out TPR domain from a symmetry mate. In *(iv)* there are no crystal contacts in the vicinity of the α_inh_.

**Figure S7.**
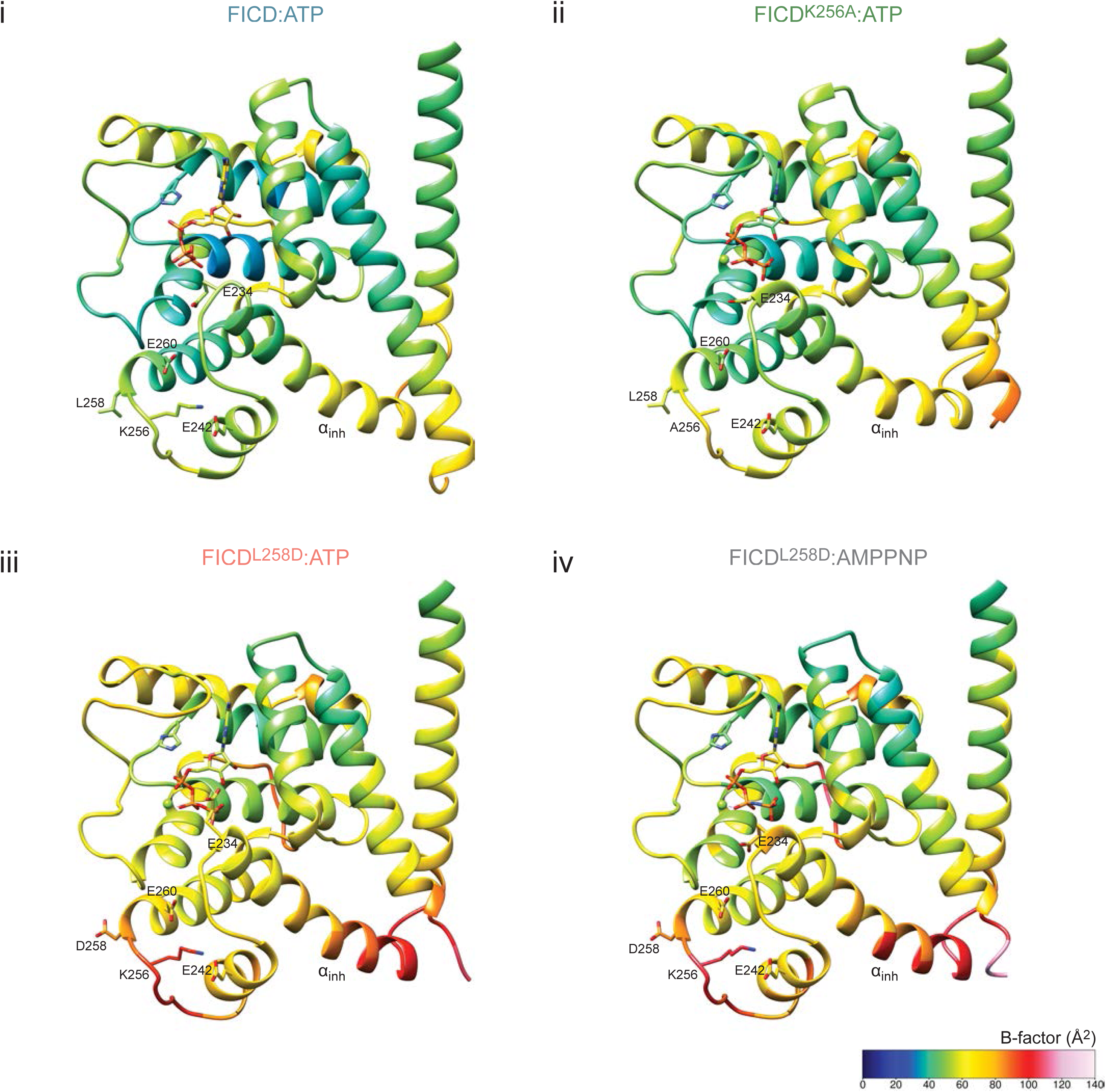
AMPylation activity correlates with enhanced flexibility of the dimer interface and Glu234. The residue average B-factors, for the four FICD complexes cocrystallised with ATP, are shown [in *(i-iv)*] with a cold to hot colour code. They display a trend of increasing B-factors in the dimer interface and in the inhibitory glutamate region. This increase in B-factor is indicative of increasing flexibility and correlates with greater AMPylation activity of the corresponding FICD. All of these structures have almost identical dimer packing in their respective crystals and limited crystal contacts around the inhibitory helix (see Figure S6). Note, structure averaged B-factors are comparable (see Table 1). For clarity, the TPR domain (up to residue 182) is not shown.

**Figure S8.**
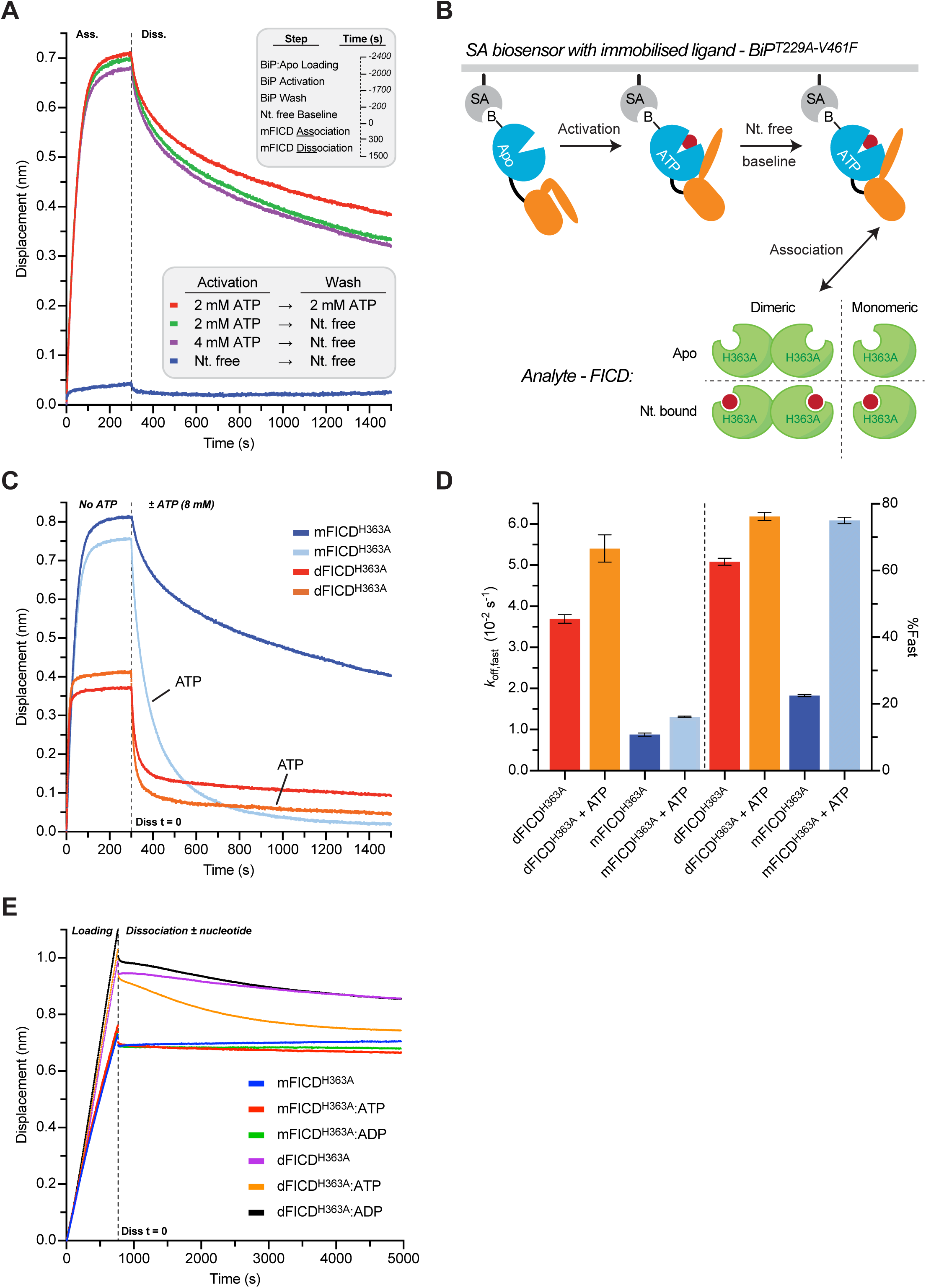
ATP negatively modulates pre-AMPylation complex and FICD dimer stability. A) Immobilised BiP responds allosterically to, is saturated by and retains ATP for the duration of BLI kinetic assays. BLI traces of the interaction between FICD^L258D-H363A^ and immobilised biotinylated BiP^T229A-V461F^ in different nucleotide states. Before exposure to FICD^L258D-H363A^ immobilised BiP:Apo was subjected to two consecutive incubation steps (activation and wash) in the presence or absence of ATP as indicated. FICD association and dissociation steps (shown) were then conducted in a nucleotide (Nt.)-free solution. Note that BiP only interacts with FICD^L258D-H363A^ when pre-saturated with ATP. Importantly, ATP pre-bound BiP retains its affinity for FICD^L258D-H363A^ even if subsequently washed in a buffer lacking ATP (compare red and green traces). Thus BiP retains its bound ATP for the duration of the kinetic experiment, experimentally uncoupling the effect of nucleotide on the FICD analyte from its effect on the immobilised BiP ligand.
B) Cartoon schematic of the BLI assays presented in Figures 6A-B. The pre-AMPylation complex is formed between the immobilised BiP:ATP ‘ligand’ and the FICD ‘analyte’.
C) The BLI association and dissociation traces from Figure 6B are shown. The immobilised biotinylated BiP^T229A-V461F^ was saturated with ATP and then exposed to nucleotide-free FICDs. Dissociation was performed in absence or presence of ATP, as indicated. [mFICD^H363A^: FICD^L258D-H363A^; dFICD^H363A^: FICD^H363A^].
D) Quantification of the biphasic exponential decay fitting of dissociation traces shown in Figure 6B. Relative ATP-induced changes of these kinetic parameters are given in Figure 6D. Shown are mean values ± SD from three independent experiments. Note the greater relative contribution of fast dissociation of mFICD in presence of ATP versus absence.
E) Representative BLI traces of an FICD dimer dissociation experiment plotted in Figure 6E. The legend indicates the form of unlabelled FICD incubated with the N-terminally biotinylated FICD (at a 100-fold molar excess, prior to biosensor loading) and also the ligand present in the dissociation buffer (at 5 mM) if applicable. Note, probes loaded with biotinylated FICD incubated with mFICD^H363A^ act as controls for non-specific association and dissociation signals, these were subtracted from the respective dFICD^H363A^ traces in Figure 6E.

**Table S1.**
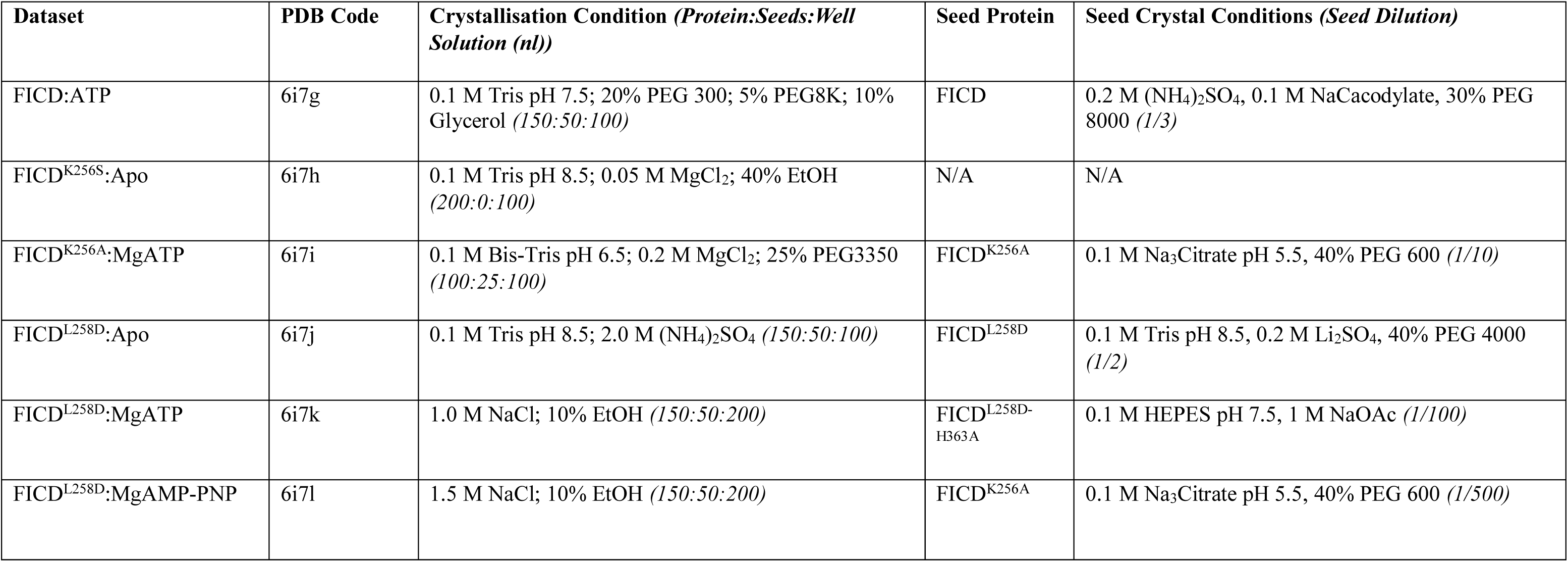
Crystallisation conditions. Where applicable the crystallisation conditions (and seed dilution) of the crystals used for micro-seeding are also shown. Note, PEG percentage is given in w/v and EtOH percentage in v/v.

**Table S2** List of plasmids used, their lab names, description, their corresponding label and references.

